# Population structure, genetic diversity and global migration of the banana Fusarium wilt pathogen, *Fusarium oxysporum* f. sp. *cubense* tropical race 4

**DOI:** 10.64898/2026.07.23.740282

**Authors:** Emmanuel Wicker, Megan Ceris Matthews, Sebastien Ravel, Jaime Aguayo, Sandrine Fabre, Nadia Adjanoh-Lubin, Céline Lopez-Roques, Jules Sabban, Joanna Lledo, Thangavelu Raman, Wayne O’Neill, Saif M. Alkaabi, Maged Elkakhy, Chunyu Li, Siwen Liu, Ganyun Yi, Jamisse Amisse, Beatrix Coetzee, Altus Viljoen, Diane Mostert

**Author notes:** **Author for correspondence**: Emmanuel Wicker, Diane Mostert.

## Abstract

- The current epidemic of banana Fusarium wilt is caused by *Fusarium oxysporum* f. sp. *cubense* tropical race 4 (Foc TR4). Foc TR4 originates in Asia and was first reported in the early 1990s, whereafter it was restricted to five Asian countries and the Northern Territory of Australia for two decades. Since 2013, the fungus was detected in the Middle East, the Indian subcontinent, Africa and Latin America. Its recent spread may be caused by the expansion of Cavendish banana production and the movement of contaminated planting materials.
- Whole genome sequences of 119 Foc TR4 isolates were compared based on single nucleotide polymorphisms (SNPs) to assess the population structure, genetic diversity and relation among isolates from 20 countries to reveal the evolutionary origin and likely migration.
- Foc TR4 is a monophyletic lineage divided into four clusters. Two of the clusters included isolates mostly from eastern Indonesia and seem to be ancestral with many rare alleles and high genetic diversity supporting this region as centre of origin. The remaining clusters were less diverse and associated with pandemic expansion.
- Knowledge on Foc TR4 emergence and dispersal can be used to inform more effective biosecurity and management practice.

## Introduction

Plant pathogens and their hosts share a long history of co-evolution in nature (Stukenbrock & McDonald, 2008). The transition from informal to organised agriculture, however, has upset this balance due to the planting of highly homogeneous, even clonal, crops over large areas. This resulted in selection pressure on pathogens to overcome plant disease resistance (Tellier & Brown, 2011). The development of trade between countries also favoured the migration of pathogen-contaminated plant material, as pathogens were taken from their native range to new environments where they caused disease to native hosts with which it did not share an evolutionary history (Zhan *et al*., 2015). These emerging diseases pose serious threats to global food security and jeopardise the survival of host crops.

Few biological invaders have caused as much havoc on a cultivated plant species as *Fusarium oxysporum* f. sp. *cubense* (Foc), the causal agent of Fusarium wilt of banana (Stover, 1962). Foc has been responsible for two major epidemics (Viljoen *et al*., 2020a). The first epidemic, caused by Foc race 1, resulted in significant damage to Gros Michel bananas grown for export in Central America in the early 1900s (Stover, 1962). The heavy reliance on a single variety, farmed in monoculture, coupled with the fact that plantations were established with infected suckers, resulted in severe losses. The direct financial impact of Fusarium wilt on the Gros Michel-based trade was estimated to be at least US$ 2 billion in 2000 figures (Ploetz, 2005). After Gros Michel was replaced with the Cavendish banana, the international banana trade grew rapidly. Tissue culture technology enabled the mass production of disease-free and genetically identical plants with grower-preferred traits to establish new plantations. Today, almost 50% of total global banana production consists of Cavendish banana (Lescot, 2015).

The discovery of Fusarium wilt on Cavendish bananas in Indonesia and Malaysia in the early 1990s revealed the start of the second epidemic, caused by a new strain known as Foc tropical race 4 (Foc TR4). Foc TR4 has now officially been reported from 25 countries in Asia, the Middle East, Africa, and Latin America (Viljoen *et al*., 2020a; Munhoz *et al*., 2024). The movement of infected planting material has primarily been associated with the spread of the fungus due to limited awareness and a disregard for quarantine measures (Viljoen *et al*., 2020a). It was estimated that Foc TR4 could affect 17% of global banana-production by 2040, which estimates to 36 million tons of bananas worth over US$ 10 billion (Scheerer *et al*., 2018).

It is widely believed that the Foc TR4 co-evolved with bananas in Indonesia and Malaysia (Ploetz & Pegg, 1997). Mass decline due to Foc TR4 was noticed when Cavendish monoculture plantations were first established in both countries in the early 1990s (Ploetz, 2000; Buddenhagen, 2009). After that, the fungus was also reported in mainland China (Li, CY *et al*., 2013), the Northern Territory in Australia (Conde & Pitkethley, 2001), the Philippines (Molina *et al*., 2009) and Taiwan (Hwang and Ko, 2004). Foc-TR4 remained restricted to the Asian countries and northern Australia for more than 20 years (Molina *et al*., 2009; Mostert *et al*., 2017). Secondary spread to the Middle East and Africa occurred almost simultaneously. Foc TR4 was first identified in the Middle Eastern country, the Sultanate of Oman, in 2012, but was probably present as early as 2009 (Viljoen *et al*., 2020a). After its first introduction, Foc TR4 was also reported in Lebanon and Jordan (Garcia-Bastidas *et al*., 2014), and subsequently in Pakistan (Syed *et al*., 2015), Israel (Maymon *et al*., 2018), and Türkiye (Özarslandan & Akgül, 2020). In Africa, Foc TR4 was first detected in northern Mozambique in 2013 (Viljoen *et al*., 2020b), where it was confined, until it was found in the nearby islands in the Comorian Archipelago. The pathogen was first found in Mayotte in 2019 (Aguayo *et al*., 2021) and later in the Grand Comoros Island (Mmadi *et al*., 2023).

Foc TR4 has spread rapidly to new countries in Asia from 2016. After it was reported from Pakistan in 2015 (Ordoñez *et al*., 2015), it was also detected that same year in the Bihar state of India (Thangavelu, R. *et al*., 2019). Since then, Foc TR4 has rapidly spread in Bihar and to the states of Uttar Pradesh, West Bengal, Gujarat, Maharashtra and Madhya Pradesh, with devastating consequences (Thangavelu *et al*., 2024). Foc TR4 was also introduced into Laos, Myanmar, and Vietnam (Chittarath *et al*., 2018; Hung *et al*., 2018; Zheng *et al*., 2018; Chittarath *et al*., 2022), probably due to the expansion of commercial Cavendish production from China into neighbouring countries. It is now also found in Nepal (Khadka *et al*., 2026) and Bangladesh (Islam *et al*., 2026). In Australia, Foc TR4 was detected in the country’s main banana production region, northern Queensland, in 2015 (O’Neill *et al*., 2016). This comes almost 20 years after its initial report in the Northern Territory more than 2000 km away (Pegg *et al*., 2021).

Concerns regarding the potential introduction of *Fusarium oxysporum* f. sp. *cubense* Tropical Race 4 (Foc TR4) into Latin America have long been emphasized, given the profound socio-economic reliance of the region on banana production (Loeillet & Dawson, 2023; Munhoz *et al*., 2024). These apprehensions were substantiated when Foc TR4 was officially reported in Colombia in 2019 (García-Bastidas *et al*., 2020). Since its introduction into the La Guajira region it has spread to Magdalena, a department located more than 150 km away from the initial outbreaks (Munhoz *et al*., 2024). Foc TR4 was also reported from Peru in 2021 (Acuña *et al*., 2022), and from Venezuela in 2023 (Mejias Herrera *et al*., 2023) where it has rapidly spread in banana fields for domestic consumption (Munhoz *et al*., 2024).

The demographic history and gene flow among geographic pathogen populations can provide information on the origin of the organism (Morris *et al*., 2010; Brazier *et al*., 2022). Combined approaches to estimate genetic diversity and compare population structures have been employed to construct migration scenarios for various plant pathogenic species (Goss, 2015). No teleomorph has been identified within the *F. oxysporum species* complex and Foc TR4 isolates belong to a single vegetative compatibility group (VCG) complex, VCG 01213/16, which are clonally related (Ordoñez *et al*., 2015). To infer genetic variation within populations with low levels of diversity, such as Foc TR4, whole genome single nucleotide polymorphisms (SNPs) have been commonly used (Goss, 2015). (Zheng *et al*., 2018), used SNPs to demonstrate that Foc TR4 isolates from Vietnam, Laos, and Myanmar were closely related to those in the Yunnan province of China, which suggested that its incursion and spread in the Mekong region were likely due to the movement of infected material and equipment from China (Chittarath *et al*., 2022). Foc TR4 genomes from Jordan, Israel and Lebanon were also closely related, allowing (Maymon *et al*., 2020) to propose that the pathogen most likely spread within the Middle East after a single introduction. In contrast, (Reyes-Herrera *et al*., 2023) showed that the introduction of Foc TR4 into Latin America was not a single event, and that isolates from Peru were different from those in Colombia.

To date, studies to investigate the origin of Foc TR4 incursions have been regional and limited by the small number of genomes available. In this study, a global collection of 119 Foc TR4 genomes was used to infer the population structure and genetic relation among isolates from different countries from where it was reported, to investigate the evolutionary origin and direction of migration.

## Materials and Methods

### Fungal collection

One hundred and nineteen Foc TR4 genomes were selected to represent 20 Foc TR4-affected countries, including 97 Foc TR4 (VCG 01213/16) isolates that were newly sequenced and 22 samples retrieved from the National Centre for Biotechnology Information (NCBI) (Fig.1). The cultures were collected between 1990 and 2019 from different locations and a variety of banana cultivars (Table S1). Localisation, sampling conditions, and storage conditions are detailed in Methods S1 and Table S1. Isolates were previously confirmed as Foc TR4 with molecular assays (Li, MH *et al*., 2013; Aguayo *et al*., 2017; Matthews *et al*., 2020) and VCG testing (Bentley *et al*., 1998; Molina *et al*., 2009; Li *et al*., 2011; O’Neill *et al*., 2016; Mostert *et al*., 2017; Chittarath *et al*., 2018; Hung *et al*., 2018; Thangavelu, Raman *et al*., 2019; Viljoen *et al*., 2020b; Aguayo *et al*., 2021; Chittarath *et al*., 2022).

**Figure 1.**
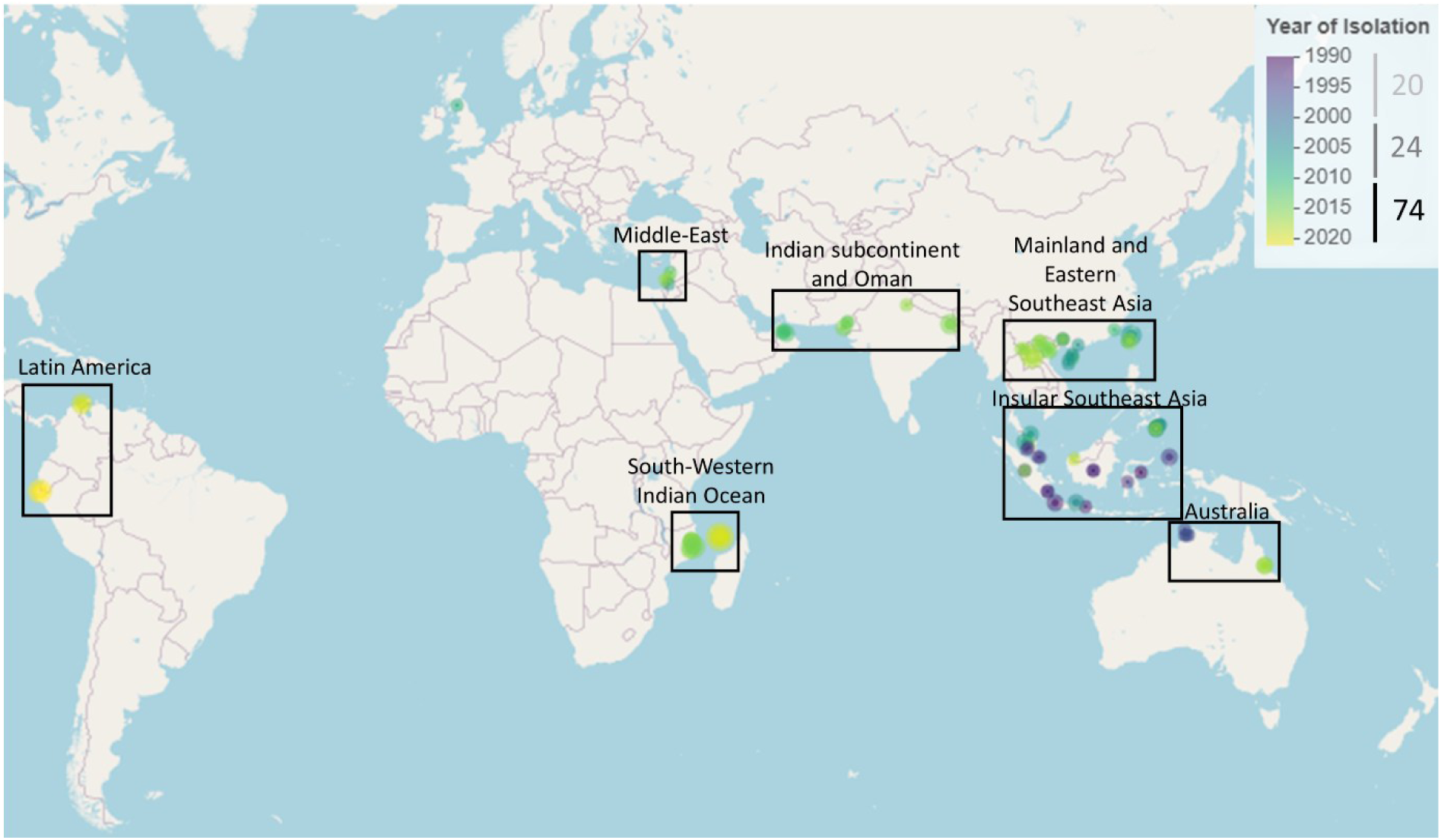
The Foc TR4 collection. A total of 119 genomes of Foc TR4 collected between 1990 and 2021 were studied, including 22 retrieved from NCBI. Sample locations were coloured according to their year of isolation. The collection was sampled from 20 countries and seven geographical regions determined from GPS coordinates (boxes): from East to West and South to North, 1. Australia, 2. Insular-Southeast-Asia (Indonesia, Malaysia, Philippines), 3. Mainland and Eastern Southeast Asia (China, Laos, Myanmar, Taiwan, Vietnam), 4. Indian Subcontinent and Oman (India, Pakistan, Oman), 5. South-Western Indian Ocean (Mayotte, Mozambique), 6. Middle-East (Israel, Jordan, Lebanon), 7. Latin America (Colombia and Peru). The UK genome clustered geographically with the Middle-East genomes, but was considered separately.

### DNA extractions, library preparation and genome sequencing

DNA of the newly sequenced isolates were extracted using the Qiagen DNeasy Plant Mini Kit (Qiagen, Hilden, Germany) with adaptations, and a MATAB protocol (Methods S1). DNA integrity was assessed with gel electrophoresis, while its quantity and quality were determined by spectrometry. The Foc TR4 isolates were sequenced on an Illumina platform at Novogene in Singapore (https://jp.novogene.com) and Get-PlaGe in Toulouse, France (https://get.genotoul.fr/) (Table S1, Methods S1).

### Sequence analysis and variant calling

A Snakemake pipeline developed in-house at CIRAD, Montpellier, called “rattleSNP” (https://forge.ird.fr/phim/sravel/RattleSNP, https://rattlesnp.readthedocs.io/en/latest/), was used to automate quality checks, Illumina read trimming, alignment to the reference genome, SNP calling, variant call format (VCF) filtering, and data conversion for downstream phylogeny and population genomics analyses (Methods S1). The output data files include multifasta and vcf files (Fig. S1). Quality control, reference mapping to the Foc TR4 reference genome UK0001 (Warmington *et al*., 2019), and variant filtering were performed as described in Methods S1. For downstream analyses, only SNPs (excluding indels) with a minimum read depth of 8, a minimum Phred quality score of 30, and no missing data were retained.

### Phylogenomic analysis

In a preliminary analysis, whole-genome SNPs were called on Foc TR4 (n=119) and a selection of other Foc VCGs, such as VCGs 0120/15, 0121, 01210 (n=16) (Table S1). A Maximum-Likelihood (ML) phylogeny was built using RAxML-NG (version 1.0.1) (Kozlov *et al*., 2019) with node support scores tested over 1000 bootstraps. The among-site heterogeneity was estimated using a discrete gamma model. The number of trees to generate for random and parsimony parameters was set to 50. The ML tree was then visualised and annotated using FigTree v1.4.4. (http://tree.bio.ed.ac.uk/software/figtree/) and iTOL version 6.5.7 (https://itol.embl.de/). File conversion and manipulation were done using R packages vcfR (v.1.12.0 and 1.15.0) (Knaus & Grunwald, 2017), adegenet (version 2.1.7) (Jombart, 2008) and DartR (v. 1.9.9.1 and 2.9.7) (Gruber *et al*., 2018; Mijangos *et al*., 2022). Phylogenetic networks were computed using the neighbor-net method implemented in SplitsTree4 (Huson & Bryant, 2006) to test for signatures of past recombination events.

### Analysis of the population structure

The population structure of the global Foc TR4 SNP data set was investigated with a combination of SnapClust and DAPC to estimate the most probable number of population clusters. SnapClust combines an expectation-maximisation (EM) algorithm with geometric approaches (Beugin *et al*., 2018). Ward’s clustering (Murtagh & Legendre, 2014) was used to define the initial clusters, and the method calculated the likelihood of each clustering solution for a varying number of clusters, k. The Akaike Information Criterion helped to choose the optimal number of clusters in the dataset when the maximum number of clusters was set to 20. The expectation-maximisation (EM) algorithm was run for 100 iterations. The optimal clustering solution and k were then used as prior-to-run DAPC, which maximises variance between groups while minimising variance within groups. For DAPC, 50 principal components were retained, and posterior group assignments made for each Foc TR4 genome as a measure of the admixture proportion originating in each cluster. Foc TR4 genomes were assigned to the clusters with the highest posterior group assignments. Clusters identified were named TR4-SC throughout the text.

### Population genetic diversity and divergence, neutrality tests

Populations were defined as country populations and TR4-SC populations (detailed in Table S2). Samples from Myanmar, Lebanon, the UK, Jordan, Israel were excluded from the population diversity analysis because of low sample sizes.

### Multi-locus genotypes and lineages (genotypic diversity)

To measure the genotypic diversity, the number of multi-locus genotypes (MLGs) and lineages (MLLs) per population were assessed first. Multi-locus lineages (MLLs) were defined from MLGs using the genotyping error as threshold (Shakya *et al*., 2021): the genotypes differing only within the genotyping error were collapsed into a same MLL, to remove the effect of technical error and groups variation. To determine the genotyping error threshold, the Euclidean distance between two CAV 807 genomes sequenced on an Illumina platform was calculated on the SNP dataset. The distance between the technical repeats (8.602) was then set as threshold and MLGs were collapsed into MLLs. A rarefaction curve was constructed to compare the MLL diversity per country population and SC clusters, considering sample size differences, by using the function *rarecurv* of the R package vegan v.2.4-2 (Oksanen *et al*., 2024).

#### Nucleotide diversity

The genome-wide nucleotide diversity π (average number of pairwise differences) per site was calculated (Hohenlohe *et al*., 2010) and expressed as a mean nucleotide diversity for each country and TR4-SC cluster, using R v.4.3.3. and the package snpR v.1.2.9.1. (Hemstrom & Jones, 2023).

### Tajima’s D neutrality test

Tajima’s test was used to identify sequences that did not fit the neutral theory model at equilibrium between mutation and genetic drift. Tajima’s D (Tajima, 1989) compares estimates of diversity θ based on either the number of observed pairwise differences (Tajima’s θ) or the number of segregating sites (Watterson’s θ). Since low frequency minor variants contribute to these statistics, it relies on the ratio of the number of variants vs the number of sequenced non-polymorphic sites. The test was performed on unfiltered SNPs (no filtering on the minimum allele frequency using the calc_tajimas_d function of the R package snpR v.1.2.9.2 (Hemstrom & Jones, 2023), on non-overlapping sliding windows of 100 kb.

### Genetic differentiation

To assess whether the Foc TR4 dataset was structured by geography, and to trace putative migration patterns, the genetic differentiation among country populations were assessed. The total genetic variance in a country population, relative to the total genetic variance of the global population (F_st_), can be used to infer differentiation between populations (Wright, 1965; Meirmans & Hedrick, 2011). The fixation index (F_st_) was calculated between each pair of populations based on SNP markers. The range of F_st_ is from 0 to 1, with 0 indicating no differentiation and high gene flow, and 1 indicating a high differentiation and no gene flow. In this study, F_st_ was calculated based on the formula of (Weir & Cockerham, 1984) with the function *gl.Fst.pop* of the R package DartR. Significance of pairwise FST was tested on 1000 bootstrap replicates (P<0.05).

### Analysis of migrations

To infer the most likely origin of Foc TR4, a reconstruction of ancestral discrete states were initially performed in which the discrete states considered were the countries of origin of selected genomes in the dataset (Shakya *et al*., 2021). The whole analysis process, based on a Maximum Likelihood (ML) phylogenetic tree, is detailed in Methods S1. A maximum Likelihood (ML) tree with 1000 bootstrap replicates was constructed and a Lewis correction applied to correct for ascertainment bias using RAxML-NG. This ML tree was then used to infer the likelihood of the probability of country origin for each node.

The historical country population relationships were investigated using TreeMix software by estimating a ML tree, the amount of genetic drift in each population, and the number of migration events that best fitted the data (Pickrell & Pritchard, 2012). The first dataset consisted of 17 country populations (n ≥ 3), whereas genomes from Israel, Jordan and Lebanon were grouped (Middle East). CAV180 was used as the outgroup. The treemix input file was generated from the R::adegenet genlight dataset re-assembling SNPs and country populations, using the gl2treemix function in the R package DartR v. 2.7.2. (Mijangos *et al*., 2022). The data was analysed using the pipeline https://github.com/carolindahms/TreeMix (Dahms, 2021), which consists of a combination of bash and R scripts (Milanesi *et al*., 2017; Zecca *et al*., 2020) (Methods S1).

### Clonality and reproductive mode

#### Mapping reads to the loci MAT-1 and MAT-2

To determine the mating type of the Foc TR4 collection, Illumina reads of all genomes were mapped on the MAT-1 sequences of *Fusarium oxysporum* f. sp*. lycopersici* (GenBank AB011379.2) and Foc TR4 genome II5 (GenBank XM_031212725.1), and the MAT-2 locus of *Fusarium oxysporum* f. sp. *radicis-lycopersici* (GenBank AB011378.1). The read sequences were aligned on the reference MAT loci using the Burrows-Wheeler aligner (Li & Durbin, 2009; Li & Durbin, 2010), implemented as BWA-MEM in the RattleSNP pipeline.

#### Simulated Index of Association

The Index of Association (I_A_) (Maynard Smith *et al*., 1993; Agapow & Burt, 2001) was calculated on clone-corrected data for country populations and TR4-SC populations, using the R package poppr v. 2.9.6 (Kamvar *et al*., 2014; Kamvar *et al*., 2015). The distribution of I_A_ were simulated under clonal (90 to 100% linked SNPs), mostly-clonal (75% linked SNPs), partially-clonal (50% linked SNPs) and sexual (0 to 10% linked SNPs) modes of reproduction (Shakya *et al*., 2021) using the R package adegenet v.2.1.10 (Jombart, 2008; Jombart & Ahmed, 2011). Then observed I_A_ were compared with simulated I_A_ (Tabima *et al*., 2018), calculated on 100 replicates of 500 randomly sampled SNPs. Means of I_A_ were compared for significance using a Kruskal-Wallis Rank Sum Test (kruskal.test function of the R package agricolae v. 1.3.7 (de Mendiburu, 2023).

Several simulations with varying sample sizes (5 to 50 individuals, 200 to 1000 SNPs sampled for calculations of I_a_) were tested to consider the impact of sample sizes and SNP numbers on the resulting I_A_. The percentage of linked loci was also considered, as numbers of individuals and number of clone-corrected SNP loci varied greatly across TR4-SC clusters. I_A_ estimations were highly variable when the number of SNPs sampled was below 500. Based on these preliminary tests, we chose to calculate the distribution of I_A_ based on 100 random samplings of 500 clone-corrected SNPs. TR4-SC3 could not be tested due to its limited sample size.

#### LD-decay

The Linkage Disequilibrium decay (LDD) was estimated for each genetic cluster using the tool PopLDdecay (Zhang *et al*., 2019) with default parameters. Each population dataset was converted to a pseudo-diploidised VCF file restricted to the core genome (i.e. excluding the contigs VMNF01000013 and VMNF01000014) and further used as input. Raw LD (estimated by the parameter r²) decay curves were fitted using the Hill and Weir model, which uses the number of SNP pairs per each distance as weights to calculate the best fitted adjustment between r^2^ and physical distance. Downstream fittings and plots were completed using the R packages tidyverse (Wickham *et al*., 2019) and ggplot2 (Wickham, 2016). Following (Nieuwenhuis & James, 2016) who reviewed methods for estimating frequency of sex in fungi, the LD decay is a good estimator of the relative importance of recombination in a fungal life cycle. More specifically, “the physical distance corresponding to the LD half-decay (LD50) track well the standard interpretation of sexual frequency across fungal species”. We thus considered this LD50 distance to compare LDD curves and to infer clonality. All bioinformatics analyses were performed on the High-Performance Computing (HPC) Core Cluster of the Institut Français de Bioinformatique (IFB Core, Orsay IDRIS, France) (ANR-11-INBS-0013).

## Results

### Generation of a population genomic dataset

The average read coverage of 119 Foc genomes included in this study was 22.56, (min=7.94, max= 95.95). The global dataset filtered for no indels, all allele frequencies and no missing values, (-miss 1), thus consisted of 65 592 high-quality SNP loci (Table S3). From recent genomics studies (van Westerhoven *et al*., 2024; Zhang *et al*., 2024) the UK0001 contigs were sorted according to synteny to the reference II-5 chromosomes (Table S4) and named Ctg 01 to 15 instead of VMNF01000001 to VMNF01000015 in the following sections to ease readability. Foc TR4 filtered SNPs were located on 11 of the 15 contigs of the reference genome UK0001 (Table S4 and Fig. S2), with relatively uniform densities across contigs (13 to 18 SNPs per 10 kb) except on VMNF01000005 where density was 8.4. However, there were very few SNPs (n=65) on the accessory contig VMNF01000013. No SNPs wereidentified on the core-contig VMNF0100015.

### The Foc TR4 populations are structured into four clusters, of which two are panglobal

Phylogenetic analysis confirmed Foc TR4 isolates to be monophyletic, with high bootstrap support (Fig. S3). The genome CAV 180 (VCG 0121), was closely related though in a different cluster. Genomes of Foc isolates included from the VCGs 0120/15 and VCG 01210 were clustered each on a distinct branch. The ML tree topology clearly separated Clade A/Foc-SC1 from Clade A/Foc-SC2 (Mostert *et al*., 2022).

Clustering analyses supported four clusters within Foc TR4 (Fig. 2a; Fig. S4). These clusters, named TR4-SC1 to 4, appeared to be well structured, with very few mixed genomes (Fig. 2b). TR4-SC3 and TR4-SC4 were highly differentiated from each other, while TR4-SC1 and TR4-SC2 were less differentiated from each other, although the difference was statistically significant (Fig. 2c). From Nei distances (Table S5), TR4-SC4 was the most divergent group (Dist= 9.63-9.90 x10^-3^), followed by TR4-SC3 (Dist= 3.428-3.472 x10^-3^), whereas TR4-SC1 and TR4-SC2 were much less distant of each other (0.341 x10^-3^). Within the phylogenetic network (Fig. 2e), many reticulations were present on these two clusters, indicating phylogenetic conflicts caused by homoplasy indicating past events of genetic exchange. These four clusters differed in geographic distributions (Fig. 2d) and dates of isolation (Fig. 2f). TR4-SC3 and TR4-SC4 were mostly found in Indonesia and only collected during the 1990s, except for one TR4-SC3 isolate collected in China in 2005. TR4-SC1 and TR4-SC2 were isolated within all three year ranges (1990s, 2000s, 2010s) and were panglobal.

**Figure 2.**
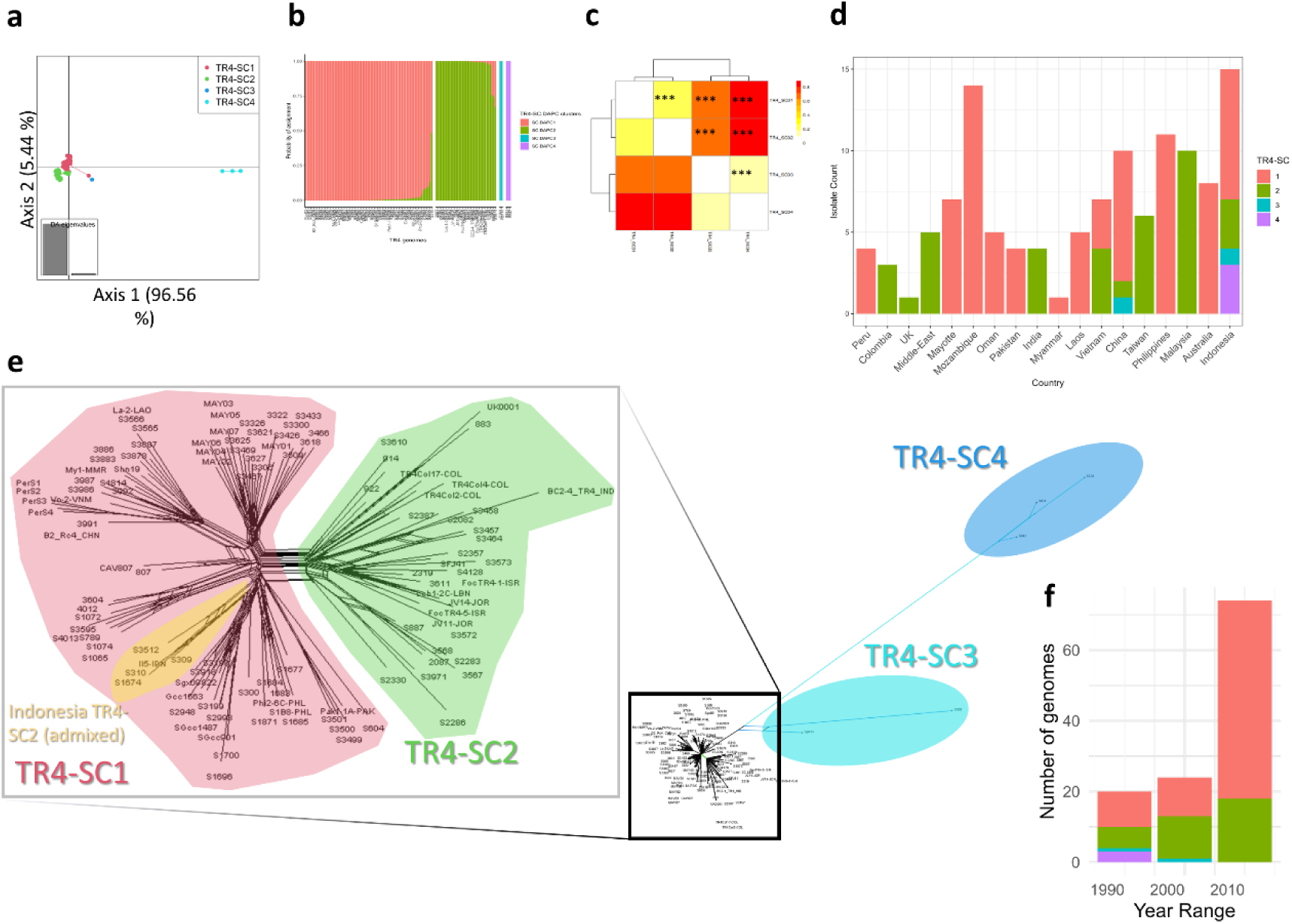
Foc-TR4 lineage is structured into four significantly differentiated subclusters, of contrasting geographical distributions and collection year ranges. (**a)** Results of Discriminant Analysis of Principal Components (DAPC) on the four clusters identified by *snapclust*. The scatterplot reports the discrimination of the four clusters (hereafter referred to as TR4-SC) based on the first and second discriminant functions of DAPC analysis. **(b)** Posterior probabilities of assignment of each TR4 genome to each of the four clusters. Each cluster looks well differentiated, with very little multiple assignments. **(c)** The four clusters are significantly differentiated (Weir and Cockerham’s Fst, 1000 bootstraps. ***: P<0.0001). **(d)** Distribution of the four TR4-SC within the countries sampled (presented from West to East). **(e)** Neighbor-Net phylogenetic network generated by SplitTree 4, showing relationships between haplotypes identified based on the full set of 65,592 SNPs without missing data. The complete dataset is presented on the right, with a zoom (left) on the main network corresponding to TR4-SC1 and TR4-SC2. **(f)** TR4-SC3 and TR4-SC4 were sampled only in the 1990s, while TR4-SC1 and TR4-SC2 were sampled throughout the three year ranges considered.

Most countries sampled appeared to be affected by isolates within only one of these two genetic clusters (Fig. 2d). The only exceptions being Indonesia (all clusters), China (TR4-SC1, TR4SC2, TR4-SC3) and Vietnam (TR4-SC1 and 2). TR4-SC1 included 77 Foc TR4 isolates with the most widespread distribution, isolated from Australia, Indonesia, Philippines, China, Laos, Vietnam, Myanmar, Pakistan, Oman, Mayotte, Mozambique and Peru. TR4-SC2 included a total of 37 Foc TR4 isolates from China, Vietnam, Malaysia, Taiwan, India, Israel, Jordan, Lebanon, the UK, and Colombia. Five highly divergent isolates were clustered on basal branches of the Foc ML phylogeny and grouped separately from the rest of the Foc TR4 isolates (Fig. S5). These divergent isolates grouped into two clusters (TR4-SC3 and TR4-SC4). TR4-SC3 included one isolate (CAV 308) from southern Java, Indonesia, while TR4-SC4 consisted of three Indonesian isolates: one from Sulawesi (CAV 831) and two from Halmahera (CAV 838 and CAV 842).

The case of the Indonesian TR4-SC2 is interesting. These isolates (n=3), collected in the early 1990s are admixed but primarily assigned to TR4-SC2. Probabilities of assignment to TR4-SC2 and TR4-SC1 for CAV 309 were 75.7% and 24.3% respectively, while II-5 assignment probabilities were 74.6% SC2 / 25.4% SC1, and CAV 1674 probabilities were 67.0% SC2 versus 33.0% SC1. All three isolates were positioned on basal branches of the ML phylogeny (Fig. S5), suggesting that they could be considered evolutionary intermediates between TR4-SC1 and TR4-SC2. The different phylogenies and analyses also positioned the reference genomes within the Foc TR4 diversity. The reference UK0001, isolated in England in the Eden greenhouse (Warmington *et al*., 2019), was positioned in the TR4-SC2 and closely related to Malaysian genomes, strongly suggesting a Malaysian origin. The reference II5, also named NRRL54006 (van Westerhoven *et al*., 2024; Zhang *et al*., 2024), was positioned in TR4-SC2 with a 74.6% probability, while assigned to TR4-SC1 at 25.4%.

### The four Foc TR4 clusters differ in diversity, demographic history and clonality

#### Genetic diversity

Of the four Foc TR4 clusters, TR4-SC4 had the highest mean nucleotide diversity (22 x 10^-3^), followed by TR4-SC3 (7 x 10^-3^), while TR4-SC1 and TR4-SC2 had very similar values that were significantly lower (both π around 0.8 x 10^-3^ (P<0.0001)) (Fig. 3a).

**Figure 3.**
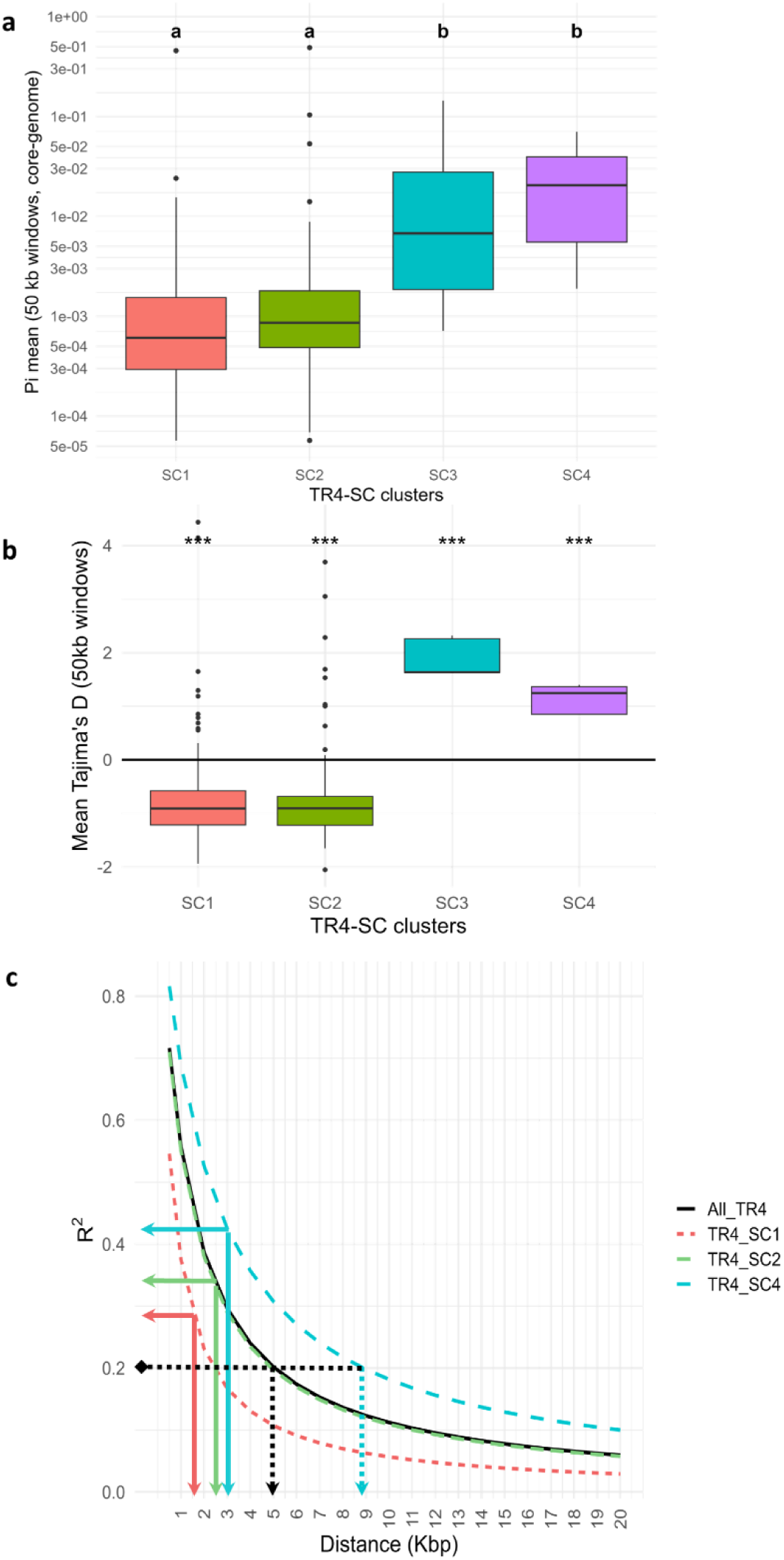
The four TR4-SC clusters differ in genetic diversity, demographic history, and clonality. **(a)** Mean nucleotide diversity π, calculated on non-overlapping 50kb windows on core-genome SNPs (without VMNF13 and 14) using the R package snpR and function calc_pi(). **(b)** Mean Tajima’s D calculated over overlapping 50 kb windows on the whole genome. All means are significantly different from 0 (Wilcoxon test, P<0.001). **(c)** Inference of the reproductive mode of the TR4-SC clusters, estimated from Linkage Disequilibrium Decay (LDD) curves. LD values (estimated by mean r^2^) were calculated on bins of 500 bp, then of 1000 bp for distances above 1000 bp. Raw LDD curves were then fitted with the Hill & Weir method, using the number of SNPpairs as weights for the model. All models successfully converged and were highly significant. Solid arrows: LDD expressed as LD50^1^= (maxLD)^-2^ (Nieuwenhuis & James, 2016). Dashed arrows: LDD expressed as LD _r=0.2_.

#### Tajima’s D

To infer demographic history of the different TR4-SC clusters, we compared values of Tajimás D reflecting the relationship between the observed and expected allele frequencies (Fig. 3b). TR4-SC3 and TR4-SC4 displayed a positive D, indicating an excess of intermediate-frequency alleles. Since positive D were observed globally on the genome (Fig. S6), this suggests that these two clusters experienced a population contraction or a population subdivision. Meanwhile, Tajima’s D observed on the panglobal TR4-SC1 and TR4-SC2 were negative (Fig. 3b), with small variations across the contigs (Fig. S6), indicating an excess of rare alleles on many loci along the genome. These values strongly suggested that TR4-SC1 and TR4-SC2 have experienced a population expansion. Collectively, these results indicated that the four SC clusters have contrasting diversities and demographic histories.

#### Reproductive mode

A single mating type, MAT-1, was identified in all Foc TR4 isolates, indicating that the fungus is asexual and suggesting a clonal reproduction mode (Table S6). The PHI and the I_A_ tests confirmed clonal reproduction when tested on the complete Foc TR4 dataset (Table S7, Figure S7). When PHI and I_a_ analysis were conducted on TR4-SCs, conflicting results were observed. From the PHI test, all clusters were recombinant (Φ=0.298, 0.294, 0.023, respectively; P-value <0.0001) (Table S7). Based on multi-locus LD (I_A_), TR4-SC4 was clonal but TR4-SC1 and TR4-SC2 were semi-clonal (Fig. S7).

LD decay (LDD) was observed for the entire Foc TR4 dataset, as well as for TR4-SC1, TR4-SC2 and TR4-SC4 (Fig. 3c); TR4-SC3 LDD could not be considered, due to its too low sample size. The whole Foc TR4 dataset displayed a LD50 distance of 2.5 kb which was similar to the one observed on TR4-SC2. The TR4-SC4 LDD curve had the least steep slope, and a LD50 of 3 kb, thus looking the most clonal. Conversely, the TR4-SC1 LDD curve slope was very steep, reaching rapidly a baseline near r²= 0.05, while the LD50 distance was 1.5 kb.

Collectively, these results strongly suggested that Foc TR4 has a semi-clonal, intermediate reproductive mode, following the terminology of (Nieuwenhuis & James, 2016). TR4-SC4 is the most clonal lineage, while TR4-SC2 and TR4-SC1 were semi-clonal, TR4-SC1 being less clonal than TR4-SC2.This may suggest that TR4 had an ancestral clonal structure (whole TR4), and that TR4-SC01 and TR4-SC02 may have experienced recent recombination events; which is consistent with the reticulations observed on the phylogenetic network.

### Analysis of country populations reveal the centre of diversity of Foc TR4

Genetic diversity was the greatest in the Indonesia population, followed by other Southeast-Asian countries such as Malaysia, Philippines, mainland China, Vietnam and Taiwan, according to rarefaction curves (Fig. 4a). Country populations outside of Asia, where Foc TR4 was more recently reported (Mozambique, Oman, Mayotte, India, Pakistan and Peru), were less diverse. The Australian population, surprisingly, had the lowest genetic diversity while Foc TR4 isolates have been collected from 1997 to 2017 (Tables S1, S2). Comparisons of mean nucleotide diversities (π) gave similar results: highest diversity in Indonesia, followed by Malaysia, China and Vietnam, then Philippines (Fig. 4b). Oman, the first country where Foc TR4 was found outside the Asia-Pacific region was next, followed by Laos, Mayotte, Mozambique, India and Colombia, Pakistan, Australia and Peru.

**Figure 4.**
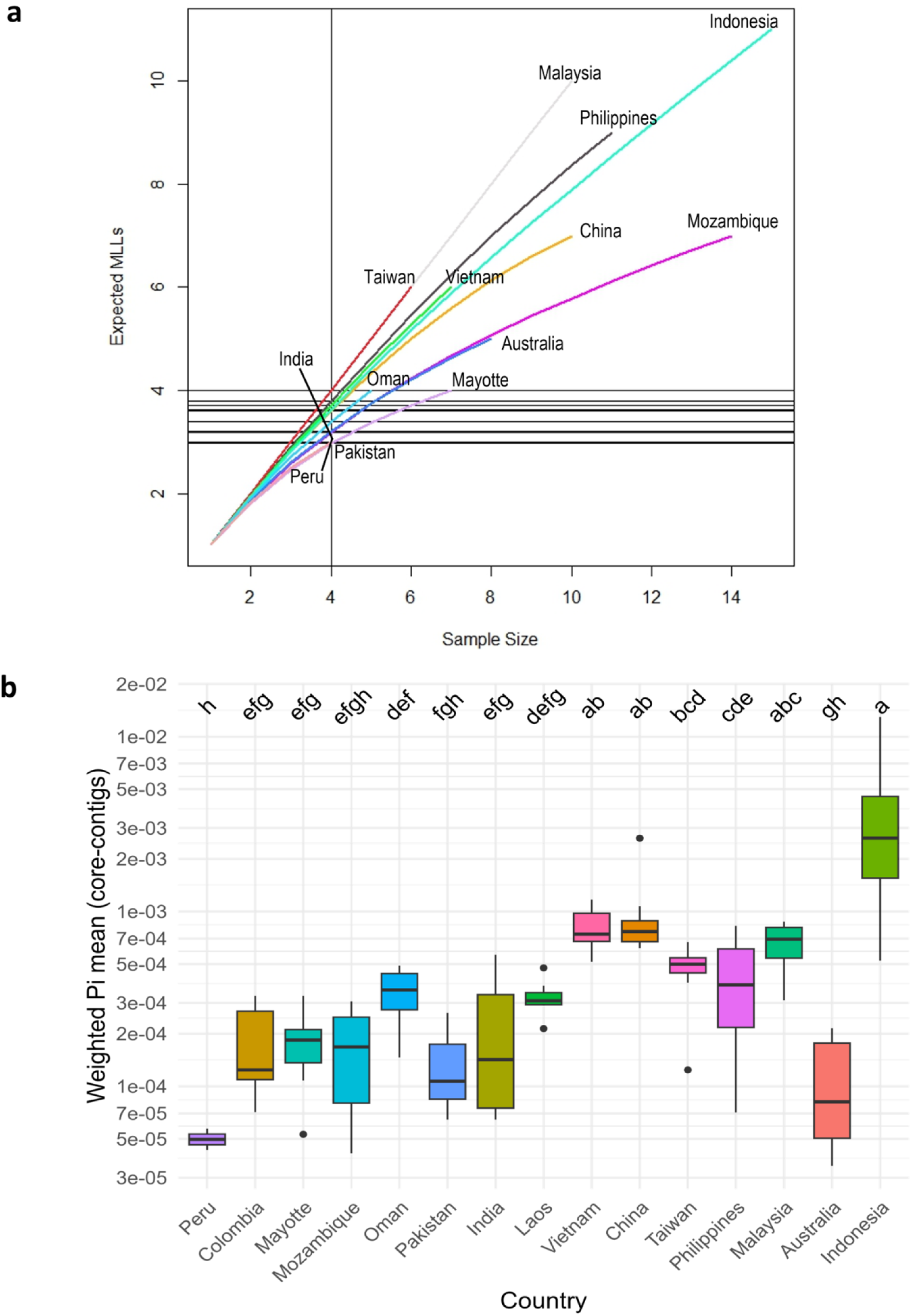
Genetic diversity is differentially distributed across countries. **(a)** Rarefaction curves of genotypic diversity, as assessed by the number of expected multi-locus lineages (MLL), across country populations, calculated using the R packages poppr and vegan. **(b)** Weighted mean nucleotide diversity π over core-contigs (i.e. excluding Contigs 13 and 14), across Country populations, sorted from West to East (Israel, Jordan, Lebanon, Myanmar, UK were not included due to their low sample sizes).

We also tested for genetic differentiation between country populations, using the pairwise Weir and Cockerham’s Fst. Under certain assumptions (migration-drift equilibrium, non-overlapping generations), Fst differentiation can be used to infer ancient and recent gene flows between populations. Patterns of genetic differentiation across countries can thus be interesting to consider, identifying putative gene flow. Indonesia had the highest number of non-significant pairwise Fsts with most other populations (Table S8). Only Australia, Philippines, Malaysia, and Mozambique, were significantly differentiated from the Indonesian population. Foc isolates from the Philippines and Oman; Mayotte and Mozambique; China, Laos, Vietnam and Peru; Malaysia, India, Taiwan and Vietnam were not differentiated, respectively. These results suggest that there was gene flow in the past within these undifferentiated populations. Collectively, these results suggest that the Foc TR4 centre of diversity may be Indonesia, with a possible secondary centre of diversification in Malaysia.

### Probable origins inferred from Ancestral State Reconstruction

To infer the ancestral relationships among country populations of Foc-TR4, a ML tree was used to infer the likelihood of the probability of country origin for each node. For a better readability, the highly divergent individuals S831, S838, S842 (TR4-SC4), and S308, Sgd13 (TR4-SC3) were removed, like was the outgroup CAV180 (Foc-SC2, VCG0121).

All basal nodes were associated with an Indonesian origin (Fig. 5), reinforcing the hypothesis of Indonesia as the unique centre of origin. Colombia and India descended from Malaysia, whereas Peru was strongly associated with a Chinese origin. Middle East (i.e. Israel, Jordan and Lebanon) was associated with a Taiwanese origin. Interestingly, the cluster including isolates from Mayotte and Mozambique was inferred to be of mixed origin, of Indonesia and Mozambique at similar likelihoods. This strongly suggests a Mozambique origin for the Mayotte outbreak. The latter Mozambique origin inferred for the common Mayotte-Mozambique node, suggests that the allele combinations found in the Mozambique populations are original and have not been observed elsewhere in our datasetAnalysis of the invasion routes and admixture

**Figure 5.**
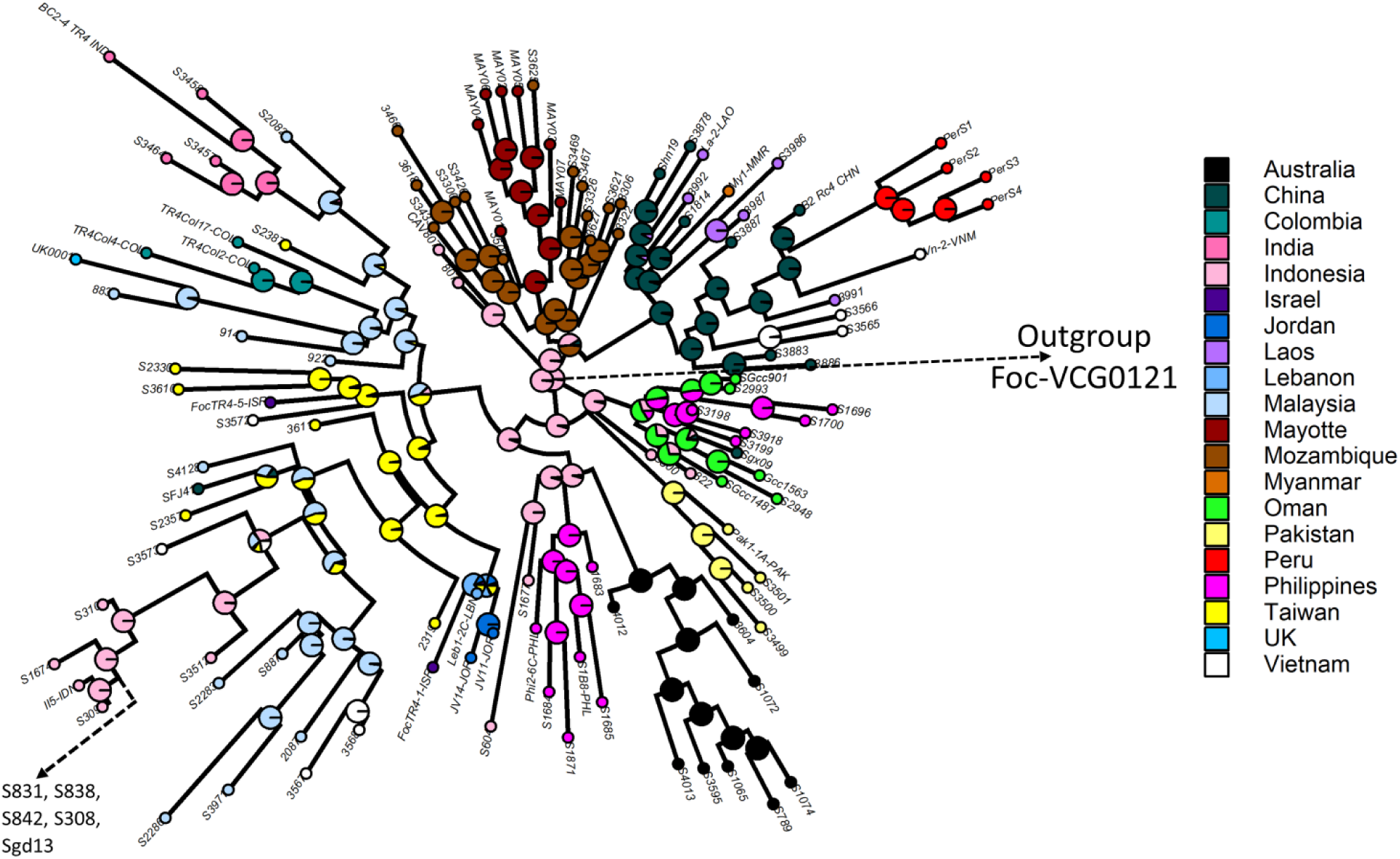
Ancestral Country reconstruction based on maximum likelihoods of nodes by geographical origin of the Foc TR4 world collection. Each label (S followed by a number) represents an individual Foc TR4 genome. The pie charts represent the likelihood of the node being assigned to each country sampled in this study.

The historical relationships among Foc TR4 country populations were estimated by modelling shared and population-specific genetic drifts among populations along a reticulated evolutionary tree, using TreeMix (Pickrell & Pritchard, 2012; Daron *et al*., 2021). The addition of sequential migration events to the initial ML tree without migration, increased the model likelihood significantly with an optimum of eight migration edges. The TreeMix tree divided into three principal branches (Fig. 6). In the first ancestral branch (inferred from the drift parameter), the populations closest to the common ancestor were Indonesia, then Philippines, up to Oman. This confirmed that Indonesia is the centre of origin and strongly suggests that the Foc TR4 in Oman originated from Philippines. The second and third branches correspond to TR4-SC1 and TR4-SC2 populations, respectively. The topology of the tree suggests that TR4-SC1 may have diverged before TR4-SC2. Moreover, the first migration edges suggested that Indonesia and Philippines both gave migrants to Malaysia and Vietnam.

**Figure 6.**
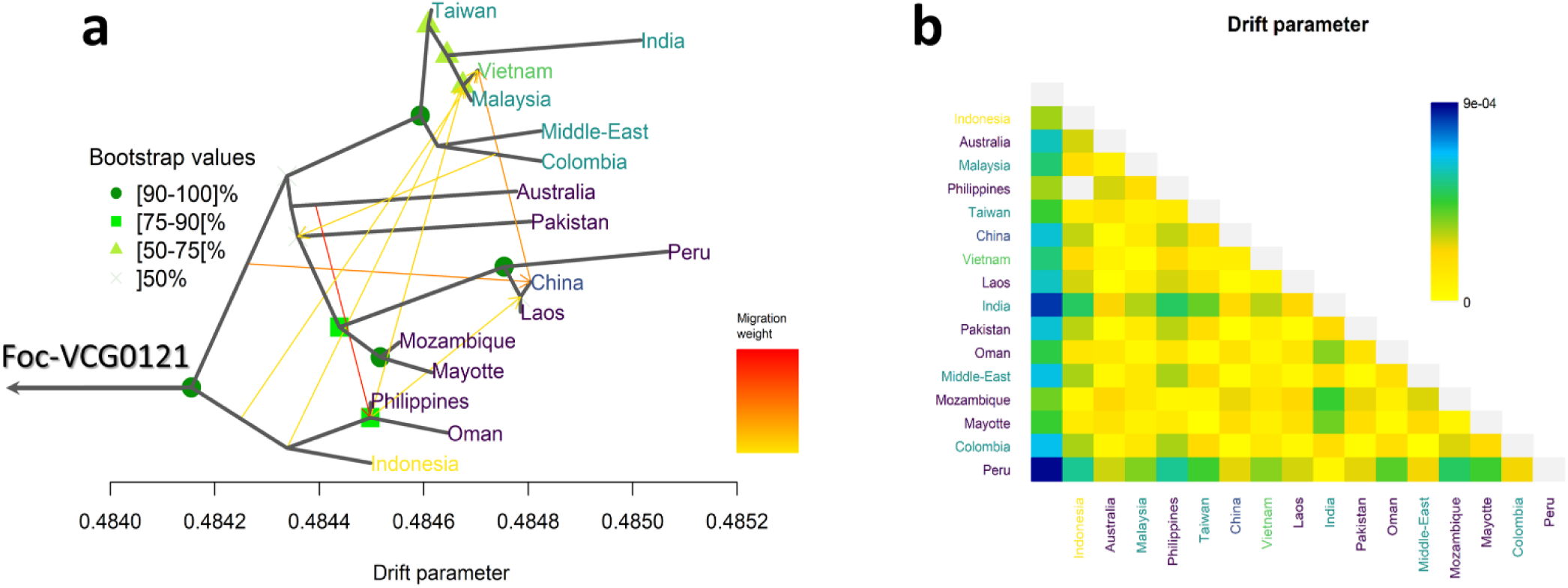
Relationships and gene flow between Foc TR4 country populations. (**a)** TreeMix ML tree with eight migration edges (arrows) rooted with CAV180 (Foc VCG 0121), including bootstrap node support. Tip colours correspond to the abundance of TR4-SC clusters: Indonesia (yellow) contains four TR4-SCs, while China contains TR4-SC1, TR4-SC2 and TR4-SC3, and Vietnam contains TR4-SC1 and TR4-SC2. TR4-SC1 countries are coloured in dark blue, while TR4-SC2 countries are coloured blue-green. (**b)** TreeMix residual matrix from the tree in (**a**).

The Australia and Pakistan populations were derived from the second branch (TR4-SC1 countries). A further subdivision separated the Mozambique-Mayotte population from the China-Laos-Peru population. Interestingly, China also received migrants from the first branch with a high migration weight, and from the Philippines. The long branches for Australia, Pakistan and Peru suggest that these country populations have undergone significant genetic drift, possibly corresponding to founder events with small population sizes and low genetic diversities. Serial founder effects during range expansion can be a spatial analogy of genetic drift (Slatkin & Excoffier, 2012).

The third branch, containing the TR4-SC2 country populations, was further subdivided into the Middle East and Colombia populations, and into a Taiwan-Vietnam-Malaysia group and Indian population. Vietnam appeared to have received introductions from China, Indonesia, and the Philippines. The topology of the tree also suggests that the Indian population probably experienced a founder event, and might have originated from Taiwan, Malaysia, or both.

## Discussion

This study is the most comprehensive phylogeographic study on Foc TR4 to date. It revealed surprising genetic diversity within Foc TR4 which were divided into four distinct clusters. It further revealed the evolutionary origin and likely migration of Foc TR4 isolates collected in Australia, Asia, the Middle East, Africa and South America. An understanding of migration patterns can help scientists and stakeholders along the banana value chain to further investigate possible introduction routes into affected countries and coordinate global efforts to enforce Foc TR4 phytosanitary measures to prevent further spread. The genomic data generated also paves the way for future pan-, comparative and functional genomics studies.

Indonesia was confirmed as the likely centre of origin of Foc TR4, as the Indonesian population was most diverse and shared ancestry with Foc TR4 isolates from most other countries included in this study. The Indo-Malayan region has previously been suggested as the centre of origin of the pathogen (Ploetz & Pegg, 1997; Fourie *et al*., 2009; Mostert *et al*., 2017; Maryani *et al*., 2019). It is believed that Foc co-evolved with its wild *Musa* hosts in the region, and that spread to new countries occurred by the movement of infected planting material (Vakili, 1965; Ploetz, 2005). Buddenhagen (2009) hypothesised that Foc TR4 (VCG 01213) most likely originated in the Malay peninsula and in Sumatra, as the pathogen was found in village banana plants soon after Cavendish banana plantations established in Sumatra and Malaya was destroyed in 1990 (Hwang & Ko, 2004; Buddenhagen, 2009). Foc TR4 isolates from eastern Indonesia, in the islands of Sulawesi and Halmahera, clustered basal to other Foc TR4 isolates in the ML tree supporting this hypothesis. An analysis of more Foc TR4 genomes from isolates collected in the 1990s or collected from native banana varieties and wild *Musa* species in the Indo-Malayan region and Pacific Ocean islands, could provide additional information and refine the area of origin of the fungus. An alternative hypothesis suggested that Foc TR4 originated in Taiwan as significant damage was reported on Cavendish as early as the 1970s (Su *et al*., 1977; Hwang & Ko, 2004). VCG testing and molecular analysis to distinguish Foc strains only became available in the late 1980s, however (Puhalla, 1985). As Foc VCG 0121, a closely related strain to Foc VCG 01213/16 able to cause disease under tropical conditions, is commonly associated with Cavendish in Taiwan (Mostert et al. 2017), it could have caused the earlier damage.

In Asia, outside the Indo-Malayan region, China and Vietnam had Foc TR4 isolates belonging to more than one TR4-SC. This suggests that the pathogen was introduced into these countries on several occasions. The population from China clustered within three TR4-SCs. Interestingly, one isolate from Guangdong where the disease was first reported in China clustered in the ancestral TR4-SC3 and was thus likely introduced from eastern Indonesia. From China, Foc TR4 was most likely moved to northern Vietnam, Myanmar (Nguyen *et al*., 2025) and Laos (Chittarath *et al*., 2018) following the expansion of commercial Cavendish cultivation. (Zheng *et al*., 2018) reported that isolates of Foc TR4 collected in the four countries were closely related, supporting the latter hypothesis. Foc TR4 is inflicting substantial damage on banana production in Laos and Vietnam and has been reported in multiple new districts since its first detection (Chittarath *et al*., 2022; Nguyen *et al*., 2025). The Foc TR4 isolates from Australia were distinct from those collected in all other countries in the region and had the lowest genetic diversity of all country populations investigated. It appears that the Australian isolates from the Northern Territory resulted from a single incursion (Conde & Pitkethley, 2001). The isolates from the Northern Territory are genetically similar to those collected in northern Queensland. Form the migration analysis, the Australian Foc TR4 isolates have probably experienced a founder event, suggesting that the introduction involved a low number of fungal genotypes.

Spread of Foc TR4 to the Middle East and Indian subcontinent were more difficult to unravel. Isolates collected in Oman grouped closely with ones from the Philippines, which suggests that an incursion is most likely from there. Spread of Foc TR4 to Lebanon, Israel and Jordan, however, is less clear. In the current study, both phylogeny and network analyses showed that isolates from these countries were most related to isolates from Taiwan. This, however, contradicts the findings of Zheng *et al* (2018) and Maymon *et al* (2020) who reported that isolates from the Middle East were closely related to isolates from the Philippines. The introduction of Foc TR4 into India is not well resolved, probably due to a low number of genomes included in the analyses. It does, however, share an ancestry with isolates from Malaysia, Taiwan or both. The isolates from Pakistan also formed a distinct population that differs from that in neighbouring countries in the region, which is likely due to a founder effect. The Pakistani isolates were closely related to those in Indonesia TR4-SC1. The two countries are known to have strong historic trade links (Malik, 2012), which might have resulted in the movement of the fungus.

It was challenging to deduce the origin of the introduction of Foc TR4 into Mozambique and Mayotte. The isolates collected in the two countries were closely related, though. Given the fact that Foc TR4 was detected in Mozambique 6 years before Mayotte, it can be hypothesised that Mozambique is the source of Foc TR4 introduction to Mayotte. The number of private alleles in Mayotte, however, suggest that Foc TR4 might have been latently present before its detection in 2019 (Aguayo *et al*., 2021). Moreover, the Mozambique populations were shown to be of mixed origin and most related to isolates from Indonesia and an unknown origin. Further samplings in this region and inferential statistics analyses may help to refine this regional scenario, taking into account the recent report of Foc TR4 in Comoros (Mmadi *et al*., 2023).

The probable origin of Foc TR4 incursions into South America were also explored. Colombian isolates clustered in a monophyletic lineage within TR4-SC2 with isolates from Malaysia; the UK genome clustered basal to this Colombian lineage suggesting it originated from Malaysia. Due to insufficient sample sizes, the exact origin of the Colombian incursion could not be determined. Maymon *et al* (2020) suggested that Colombian isolates have an Indonesian origin, but more samples will have to be included in future analysis before this theory is proven. The Peru population grouped within TR4-SC1 and were most related to isolates from China and Laos. (Reyes-Herrera *et al*., 2023) also suggested that the incursions into Colombia and Peru have been introduced from different origins. The reliance of top banana-producing countries in South America on Cavendish banana exports, and the short interval between the incursions into Colombia and Peru, are of great concern. In Peru the disease is now widespread on smallholder farms in the Piura region, where biosecurity measures are difficult to employ (M. Dita, personal communication). Countries in Latin America where Foc TR4 have not yet been reported from should thus introduce strict biosecurity practices to protect local banana industries. Awareness and regular surveillance are critical to prevent Foc TR4 from spreading in the region.

Foc TR4 is considered a clonal, asexually reproducing fungus (Fourie *et al*., 2011). Yet, the different clonality estimators used in this study suggest that Foc TR4, despite carrying a single mating type, displays varying levels of clonality. The study of (Nieuwenhuis & James, 2016) compared various fungal lifestyles and reproductive modes, and proposed a scale based on LD decay. From this scale, Foc TR4 and its subclusters showed an intermediate reproductive mode, suggesting some rare recombination events within frequent asexual multiplication cycles. The existence of a cryptic sexual state were postulated by (Taylor *et al*., 1999) and genome-wide recombination has recently been established in *F. oxysporum* f. sp. *ciceris* (Fayyaz *et al*., 2023). Mixed reproductive modes have been reported in fungi traditionally thought to be asexual such as *Coccicioides immitis*, *Aspergillus flavus* (Nieuwenhuis & James, 2016), *Macrophomina phaseolina* (Ortiz *et al*., 2023). The vascular pathogen *Verticillium dahliae* shares several similarities of life cycle with Foc. While *V. dahliae* is asexual, signatures of recombination have been found within its genome (Milgroom *et al*., 2014; Chen *et al*., 2021). Horizontal gene transfer events could, however, significantly complicate measurement of clonal structure and have shown to make populations appear more diverse than they are (Boto, 2010). Horizontal gene transfer is known to have shaped the evolution of *F. oxysporum* (Ma *et al*., 2010). In the study of (Czislowski *et al*., 2018), horizontal gene transfer was shown to influence the inheritance of the *secreted-in-xylem (SIX)* effector genes in Foc. These results warrant an extensive exploration of clonality levels within and between Foc clades.

Chromosome variations or large-scale duplications could have contributed to genetic variation observed in the Foc TR4 population. It will be important to investigate which specific regions are under selective pressure, and to confirm its associated gene contents. An early study investigating karyotype variation within Foc populations observed differences in chromosome numbers within VCGs (Boehm *et al*., 1994). These differences were believed to be driven by various factors including differences in geographic distribution range, the time that specific lineages have been genetically isolated, fitness of lineages when moved to new environments, and host specificity (Boehm *et al*., 1994). A recent pangenomic study observed large-scale duplications in *F. oxysporum* genomes and hypothesised that this plays a major role in the origin and evolution of accessory genomic regions (van Westerhoven *et al*., 2024). The acquisition of accessory chromosomes has been linked with evolution of host-specific pathogenesis in *F. oxysporum* that has been suggested for the polyphyletic nature of *formae speciales* within the *F. oxysporum* species complex (Yang *et al*., 2020). The genes associated with adaptation and chromosome arrangement and structures could, therefore, explain the lifestyle and evolution of virulence in Foc, and should be further investigated (van Westerhoven *et al*., 2024; van Westerhoven *et al*., 2025).

The phylogenomic analyses conducted in this study provide novel insights into the migration routes that have shaped the worldwide spread of Foc-TR4. TR4-SC4 and TR4-SC3, appear to be ancestral lineages due to the number of rare alleles and high genetic diversity present. Most isolates within these lineages were isolated in the early 1990s at the start of the Foc TR4 epidemic and were from eastern Indonesia. The TR4-SC2 from Indonesia is actually composed of admixed individuals isolated in the early 1990s, suggesting that TR4-SC2 and TR4-SC1 divergence occurred in Sulawesi and/or Java. The TR4-SC1 and TR4-SC2 clusters were much less diverse with less rare allele, signalling bottlenecking events and more recent adaptation after fast expansion which were associated with the more recent pan-global dispersal of the Foc TR4 epidemic. To formally estimate the probabilities of alternative invasion pathways, however, methods such as Approximate Bayesian Computation (ABC) (Guillemaud *et al*., 2010), should be considered.

The effect of changes that occur in Foc populations when sampled over temporal ranges and from different banana varieties should also be investigated in future, with more representative sampling. Integration of genomic data sampled at regular intervals during an epidemic could provide an accurate assessments of pathogen evolution and adaptability. The current study has illustrated a surprising level of genetic diversity within Foc TR4 and it highlights the importance of proper validation of molecular detection tools to encompass the diversity within the pathogen (Arrieta Salgado *et al*., 2026). It is essential to consider that sampling bias can obscure interpretation of population genomics studies and it is important to communicate new incursions with relevant national and regional phytosanitary authorities, and grower organisations, for accurate interpretations.

## Acknowledgements

Foc research and MCM stay in PHIM were funded by the CGIAR Research Program

“Roots, Tubers and Banana’ (CRP-RTB) 2020-2022, and an NRF bursary.

The ANSES Plant Health Laboratory (LSV), is supported by interdisciplinary program

ARTEMIS of Lorraine Université d’Excellence (ANR-15-IDEX-04-LUE).

The funders had no role in study design, data collection and analysis, decision to publish, or preparation of the manuscript. We thank Jean Carlier, Stéphane de Mita, Didier Tharreau, Pierre Gladieux, for useful discussions which improved the initial versions of this manuscript.

## Author Contributions

EW, AV, DM conceived and supervised the project

JAG, TR, WO, SA, ME, CL, SL, YG, JAM collected data,

MCM, NAL, SF, CLR, JS, JL, BC performed experiments

EW, SR, and MCM performed the analyses;

EW, DM and AV acquired funding for the study;

EW wrote the first draft with input from MCM, DM, and AV; all authors contributed to editing the manuscript.

## Data availability

The data that support the findings of this study are openly available in the Sequence Read Archive at https://www.ncbi.nlm.nih. gov/sra. The data sequenced in this study are available under accession no.: PRJNA1479621. The accession nos. for data from additional public projects are listed in Table S1.

## Supporting Information

**Methods S1.** Supporting materials and methods

**Figure S1.** Overview of the RattleSNP Snakemake workflow used for sequence analysis, genome calling, and variant calling.

**Figure S2.** Density of the SNPs identified within the TR4 dataset after the mapping of Illumina reads on the reference genome UK0001 (Warmington et al 2019).

**Figure S3.** Maximum-Likelihood (ML) tree generated by RaxML-NG based on whole-genome SNPs of 119 TR4 genomes and 16 STR4 genomes mapped on the TR4 reference genome UK0001.

**Figure S4.** SnapClust choose of the best K number of clusters, following the Akaike Criterion (R package *adegenet*), as applied on the global Foc TR4 genome dataset.

**Figure S5.** Maximum-Likelihood (ML) phylogenetic tree constructed with RAxML-NG v.1.0.1., using the GTR+CAT substitution model, based on the 65 592 high-quality SNPs of the Foc-TR4 dataset.

**Figure S6**. Tajima’s D calculated per TR4-SC cluster on each UK0001 contig, using calc_tajimas_d() in the R package snpR.

**Figure S7**. Inference of the reproductive mode of the Foc-TR4 whole dataset and the TR4-SC clusters, using the determination of linkage among SNP loci with the index of association (*I_A_*). **(a).** Whole-TR4 dataset. **(b)**. The TR4-SC clusters.

**Table S1.** The collection of *Fusarium oxysporum* f.sp. *cubense* tropical race 4 (Foc-TR4) isolates and genomes, and the metadata associated. The other Clade A genomes are included in the second part of the Table.

**Table S2.** Foc-TR4 populations defined by TR4-SC clusters (following SC-DAPC), sorted by geographical origins, defined as regions (clusters of GPS coordinates) and countries.

**Table S3. Read coverage statistics for the Foc TR4 dataset and the outgroup CAV180 (Foc SC2, VCG 0121).**

**Table S4. Read coverages and SNP numbers per UK0001 contig.**

**Table S5. Nei’s genetic distance (Nei 1972) between the four TR4-SC clusters (x10-3), calculated using the StAMPP::stamppNeisD function.**

**Table S6**. *In silico* determination of the mating type, by mapping the Illumina reads (BAW-MEM in the RattleSNP pipeline) on the MAT-1 gene sequences of Foc TR4 genome II5 (MAT1-TR4, GenBank XM_031212725.1) and *Fusarium oxysporum* f. sp. *lycopersici* (MAT1-Fol, GenBank AB011379.2), and the MAT-2 locus of *Fusarium oxysporum* f. sp. *radicis-lycopersici* (MAT2_Forl, GenBank AB011378.1).

**Table S7. Pairwise Homoplasy Index (PHI) tests performed on the Foc-TR4 dataset and subsets, related to TR4-SC and Country populations**.

**Table S8. Pairwise Weir and Cockerham (1984) Fst differentiation between Foc-TR4 Country populations (n≥3).**

