## Supplementary material for "Population structure, genetic diversity and global migration of the banana Fusarium wilt pathogen, *Fusarium oxysporum* f. sp. *cubense* tropical race 4": Methods S1

**Methods S1. Supporting materials and methods**

**I**

**The sample collection**

The fungal isolates, newly sequenced, originated from 13 countries in South-East Asia and Australasia (Australia, Indonesia, Malaysia, the Philippines, Laos, Vietnam), East-Asia (China, Taiwan), South Asia (India, Pakistan) West Asia (the Sultanate of Oman), and Austral Africa (Mayotte island (French overseas department), Mozambique).

Their GPS coordinates were either obtained during sample collection or determined with Google Earth (district scale). All the isolates are stored at -80℃ at the culture collection of the Department of Plant Pathology, Stellenbosch University in South Africa, or in the CIRAD Plant Health Institute of Montpellier (PHIM) in France. The *F.oxysporum* f.sp*. cubense* TR4 isolates collected for this study were collected in compliance with the Nagoya protocol (Table S1) and samples were not used for any commercial purposes.

**Geographical regions and Year-ranges**

Clusters of geographical regions were defined based on the GPS coordinates of the samples (fungal isolates and DNA sequences), following the approach presented in (Wicker *et al.*, 2012). Briefly, the physical distances between sites of collection were calculated from the GPS coordinates using R 4.3.2. (R, 2013) , the package *sp* (v.2.1-3, (Pebesma & Bivand, 2005). The obtained dataset was then submitted to a agglomerative hierarchical clustering, using the Ward criterion (R package cluster, function agnes). The best number of clusters was chosen using the hclust function, and was determined as seven Geographical Zones (Fig. 1) : (i) Insular South-East Asia : Indonesia, Malaysia, Philippines, (ii) East and Mainland South-East Asia : China, Laos, Myanmar, Taiwan, Vietnam ; (iii) Australia ; (iv) Central Asia : India, Pakistan, Oman ; (v) South-Western Indian Ocean : Mayotte, Mozambique ; (vi) Middle-East: Israel, Jordan, Lebanon ; (vii) Latin America : Colombia, Peru. The genome from UK was treated separately (see below).

Following the same hclust approach, clusters of dates of isolation were defined. Three « year-ranges » were thus identified, (i) 1990s, (ii), 2000s, (iii) 2010s.

**DNA extractions**

The Foc-TR4 isolates extracted in Stellenbosch University (Stellenbosch, South Africa) were grown in 100 mL of nutrient broth in Erlenmeyer flask with gentle agitation at 100 rpm at 25 °C in a Labcon shake incubator (Labcon, Petaluma, California, USA) for a week. The liquid cultures were then strained through cheese cloth and the mycelia of each isolate collected and transferred into 2-mL Eppendorf tubes and freeze dried in a Virtis Benchtop Pro freeze drier (SP scientific, Warminster, Pennsylvania, USA) for 48 hours. Glass beads (Company, address) were added to freeze-dried mycelia in 1.5 mL tubes and cells lysed using a Retch tissue lyser machine (Qiagen Hilden, Germany) at 30 rpm for 2 min. After, 1 mL of extraction buffer and 4 uL proteinase K at a concentration of 10mg mL-1 was added to each sample and it was mixed by inversion. The extraction buffer contained 1.4M NaCl, 0.1M Tris, 0.02M EDTA and 2% w/v cetyltrimethylammonium bromide (CTAB) and 1% w/v of polyvinylpyrrolidone (PVP). The samples then incubated in a water bath for 30 min at 60℃. After 30 min, 3 uL Rnase at a concentration of 100 mg mL-1 was added and it was returned to the water bath for another 30 minutes. Samples were then centrifuged at 12 000 x g for 10 min and the lysate transferred to another 2-mL Eppendorf tube. Buffer P3 provided in the kit was added to the lysate in a 1:3 ratio. After mixing via inversion, samples were incubated on ice for 20 mins. Samples were then centrifuged at 20 000 x g for 5 mins and the resultant supernatants transferred into QIAshredder spin columns provided in the kit. Centrifugation at 20 000 x g was repeated for 2 mins. Resultant supernatants were transferred to new 2-mL tubes and equal volumes of phenol:chloroform:isoamyl alcohol (25:24:1) (Merck, Darmstadt, Germany) added. Samples were centrifuged at 14 000 rpm for 20 mins. The top 450 uL from each sample was transferred to a new 2-mL tube. An equal volume of chloroform:isoamyl alcohol was added, samples mixed by inversion, and the centrifugation step repeated. The top 400 uL was now transferred to a new 2-mL tube and the chloroform/isoamyl alcohol step repeated. The Qiagen Dneasy extraction protocol was then followed as per the manufacturer’s guidelines from Step 6 to Step 12 with the expection of adding 100% ethanl rather than buffer AW2 provided in the kit at Step 8. In Step 11 the DNA was eluded in 30 uL elution buffer provided in the kit rather than 100 uL, and incubation occurred at 60°C rather than at 15-25°C.

DNA from the Mayotte Island isolates was extracted at CIRAD-PHIM (Montpellier, France), using a different protocol detailed below. Fungal cultures were grown for five days at 25°C in agitated liquid Nitrate medium (0.17% yeast nitrogen base, 3% sucrose, 100 mM KNO_3_ ; Mara De Sain & M. Rep, personal comm. 2020). Fungal mats were then vacuum-filtered, rinsed with sterile water, then transferred in 50 mL tubes, flash-freezed in liquid nitrogen, and stored at -80°C until use. For DNA extraction, fungal mycelium was first finely ground in liquid nitrogen, and the powder was transferred in a 5 mL tube (~1 mL powder). Lysis was first done by 2 mL of MATAB buffer (0.1M Tris-HCl pH8, 1.4M NaCl, 0.02M EDTA, 20g.L^-1^ Mixed AlkylTrimethyl Ammonium Bromide (MATAB), 10 g. L^-1^ PEG6000, 5 g. L^-1^ Na_2_SO_3_), complemented with 50μL Proteinase K (10 mg.mL^-1^). The mix was first incubated 1h at 65°C with regular twisting. Two mL Chloroform Isoamyl alcohol (CIAA) was then added to the lysate. Tubes were inverted slowly and regularly to complete the emulsion, then centrifuged for 20 min at max speed at room temperature. The supernatant was transferred in 2 mL tubes (1 mL/tube). Each tube then received 20μL RNAse A (10 mg.ml^-1^) and was incubated at 37°C for 45 min. DNA was precipitated by 900μL cold isopropanol per tube. Tubes were then softly inverted and let stand 10 min minimum in a -20°C freezer. Tubes were then centrifuged 15 min at maximum speed (~18000 rpm), and the supernatant was discarded. Pellets were washed with 1 mL Ethanol 70%, then centrifuged 5 min. Liquid was discarded by draining. Pellets were then dried 10 min in a speed-vacuum dryer, then resuspended in 110μL EB buffer (QIAGEN) for one night at 4°C.

The integrity of DNA samples was assessed with gel electrophoresis, and the quality and quantity were analysed on both a Nanodrop spectrophotometer (Thermo Fischer Scientific, Waltham, Massachusetts, United States) and with Qubit 4 fluorometer (Thermo Fischer Scientific, Waltham, Massachusetts, United States).

**Library preparation and genome sequencing**

Newly sequenced Foc TR4 isolates were Illumina sequenced at Novogene in Singapore (https://jp.novogene.com) and Get-PlaGe in Toulouse, France (https://get.genotoul.fr/), respectively (Table S1). At Novogene, DNA of each fungal isolate was sheared by sonification into fragments and sequencing libraries prepared. The quality and quantity of the DNA fragments were verified using an Agilent 2100 Bioanalyzer (Santa Clara, CA, USA). The DNA of the isolates was sequenced in four lanes of an Illumina NovaSeq6000 machine (Illumina, San Diego, CA, USA) resulting in 150 bp paired end reads. The same sequencing workflow and machine type were used in GetPlage, with sequencing performed on two lanes.

**Sequence analysis and variant calling**

*Quality control and reference mapping*

FASTQC (version 0.11.9) was used to do quality checks on raw sequence data. ATROPOS v. 1.1.31 (Didion *et al.*, 2017) was employed on Python v. 3.7 (Van Rossum & Drake Jr, 2009) to trim Illumina reads from adapter sequences. Reads were then mapped on the Foc TR4 reference genome UK0001 (Warmington *et al.*, 2019) (GenBank Number: VMNF00000000, https://www.ncbi.nlm.nih.gov/assembly/GCA_007994515.1/) using BWA-MEM (version 0.7.17) (Jung & Han, 2022). PICARDTOOLS v.2.23.5 (Broad institute. 2020) and SAMTOOLS v.1.10 (Li *et al.*, 2009) were used to tag repeated reads and evaluate genome mapping and coverage. From recent genomics studies (van Westerhoven et al., 2024; Zhang et al., 2024) (van Westerhoven *et al.*, 2024, Zhang *et al.*, 2024), we sorted the UK0001 contigs according to their synteny to the reference II-5 chromosomes (Table S4). To ease readability, we also named them Ctg 01 to 15 instead of VMNF01000001 to VMNF01000015 in the following.

*SNP calling*

SNP calling was completed using GATK4 (version 4.2.0) (McKenna *et al*., 2010) and the HaplotypeCaller function. The read mapping quality per isolate was calculated using the GATK function GenotypeGVCF, and SNP variants were retrieved using SelectVariants. VCFTOOLS (version 0.1.16) (Danecek *et al.*, 2011) was used to filter the resulting SNP markers according to sequencing depth, minimum quality, minor allele frequency (MAF) and percentage of missing data. SNP variant numbers, chromosomes covered and percentage missing data per set of parameters, were summarized in html reports. The vcf filtered datasets were compared on SNP site numbers and contigs covered, using R 4.1.1 and the package vcfR (version 1.12) (Knaus & Grunwald, 2017). Were retained for downstream analysis only the SNPs with a minimum read depth of 8, a minimum Quality (Phred Score) of 30, no missing data. The indel variants were also removed.

**Analysis of migrations**

*Ancestral area reconstruction*

To infer the most likely origins of the TR4 global populations, we performed a reconstruction of ancestral discrete states, following the approach of (Shakya *et al.*, 2021), in which the discrete states considered were the countries of origin of the genomes (tips). Our analysis pipeline was also based on the strategy presented in <https://www.phytools.org/Cordoba2017/ex/8/Anc-states-discrete.html>.

We first generated a Maximum Likelihood (ML) phylogenetic tree based on the GTR+G substitution model, using RAxML-NG v.1.0.1. (Kozlov *et al.*, 2019). The genome of CAV180 (*F.oxysporum f.sp.cubense*, clade A, cluster Foc-SC02 *sensu* (Mostert *et al.*, 2022)*,* VCG 0121) was used as an outgroup to root the tree. Based on 1000 bootstrap replicates, a ML tree was plotted using the R package ape v.5.8 (Paradis & Schliep, 2019). After removing the CAV180 root, the ML tree was used to estimate the ancestral area (or range) probabilities using the ace (ancestral character estimation) function of the R package ape. This function performs an estimation of ancestral character states for discretely valued traits using a continuous-time Markov chain model, commonly known as the Mk model method. We estimated the likelihoods of each state on the nodes following the « Equal Rate » (ER) model. In this model, transitions between states are done following symmetrical rates. The different states (countries) are thus mapped onto nodes. The empirical Bayesian posterior probabilities supporting each ancestral origin were overlain on the nodes of ML trees using the R package phytools v. 1.9-16 (Revell, 2024).

*TREEMIX analysis*

The software TreeMix was used to investigate the historical population relationships by estimating a ML tree, the amount of genetic drift in each population, and the number of migration events that best fitted the data (Pickrell & Pritchard, 2012).

The first dataset consisted of 17 country populations (n ≥ 3), where Israel, Jordan and Lebanon were grouped in Middle East. The outgroup was CAV180, as previously. The treemix input file was generated from the R::adegenet genlight dataset rassembling SNPs and populations, using the gl2treemix function in the R package DartR v. 2.7.2. (Mijangos *et al.*, 2022). We then followed the approach and pipeline developed by Carolin Dahms (<https://github.com/carolindahms/TreeMix>)(Dahms, 2021), consisting of a combination of bash and R scripts (Milanesi *et al.*, 2017, Zecca *et al.*, 2020) organized in four steps. First, a consensus ML tree was built from 500 bootstrap replicates using the PHYLIP (v.3.697) consense program. This consensus tree (containing no migration event, m=0) was then used (option -tf) to calculate the number of migration events (m) that best fitted the data by running 15 repetitions of Treemix for each m value (ranging from 1 to 15). Linkage Disequilibrium (LD) among SNPs was considered by grouping SNPs by blocks of 500 (-k 500). Secondly, the optimum number of m was identified on this « test_migrations » dataset using the R package OptM (v.0.1.8), based on two main methods, « Evanno » and « linear ». Thirdly, a consensus ML tree including bootstrap node support was obtained from 500 bootstrap runs of Treemix with the optimum number of migration edges (m), rooted on CAV180 (-root), based on the consensus tree with no migrations (-tf), blocks of 500 snps (-k 500). Then a final set of 30 Treemix runs was performed with the optimum number of migration edges (m), rooted on CAV180 (-root), based on the consensus tree with m migrations (-tf), blocks of 500 snps (-k 500). Fourth, post-processing of the final runs (comparison of tree likelihoods, ML tree with boostraps values and migration weights) were done using the R package BITE (v. 1.2.0008) with R v.3.6.3.
