## Supplementary figures and images for "Population structure, genetic diversity and global migration of the banana Fusarium wilt pathogen, *Fusarium oxysporum* f. sp. *cubense* tropical race 4"

### Fig.S1

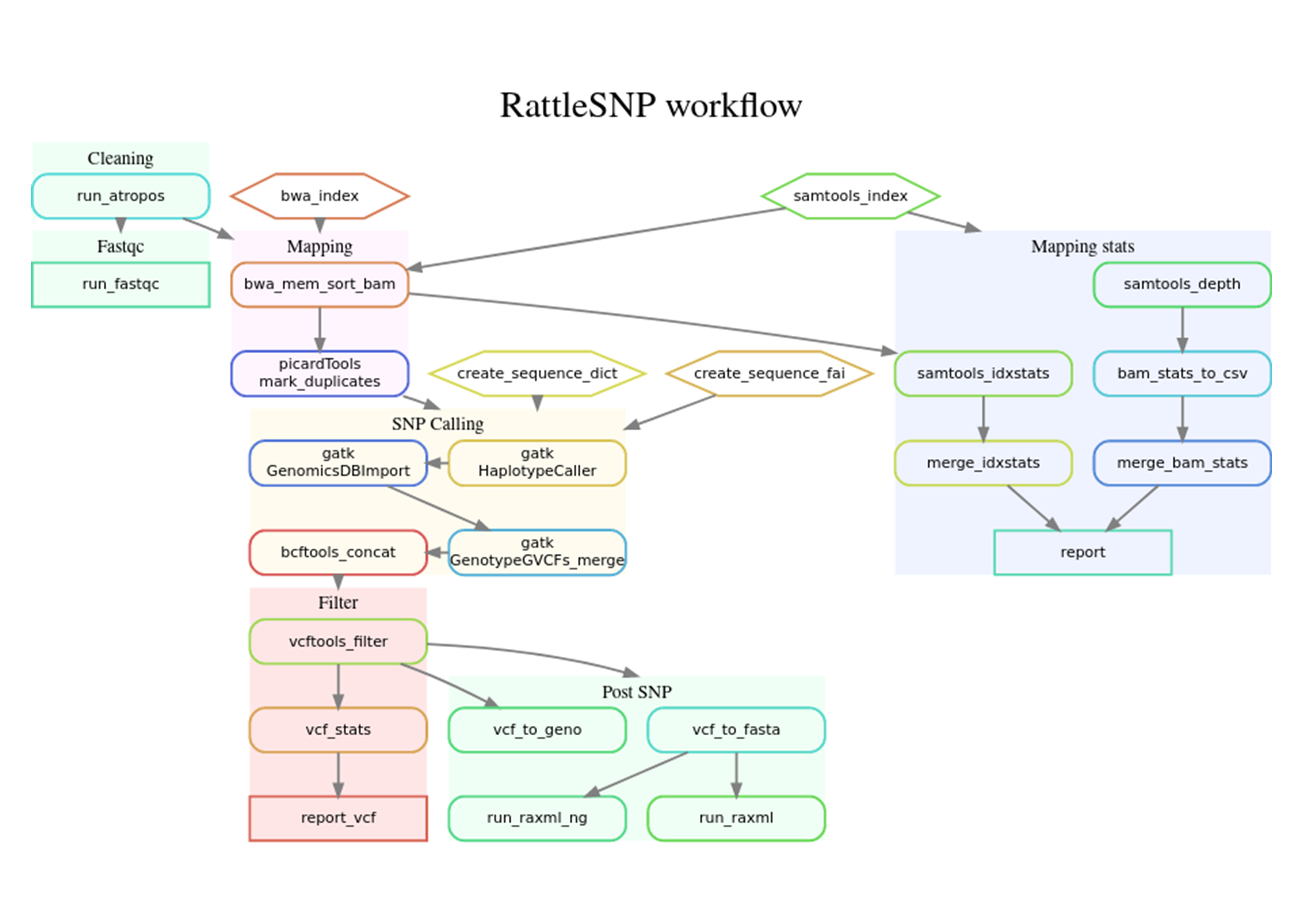

### Fig.S2

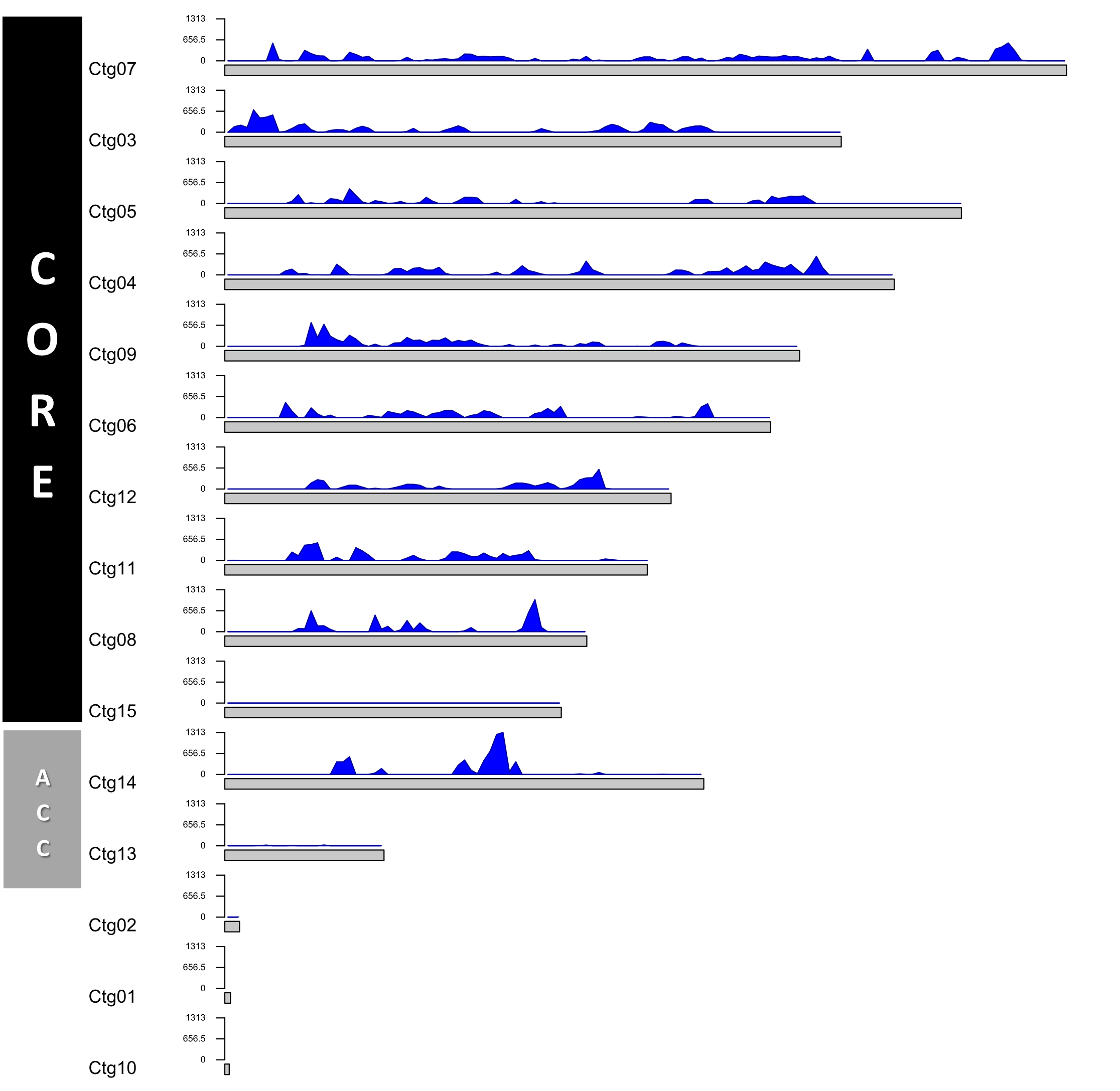

### Fig.S3

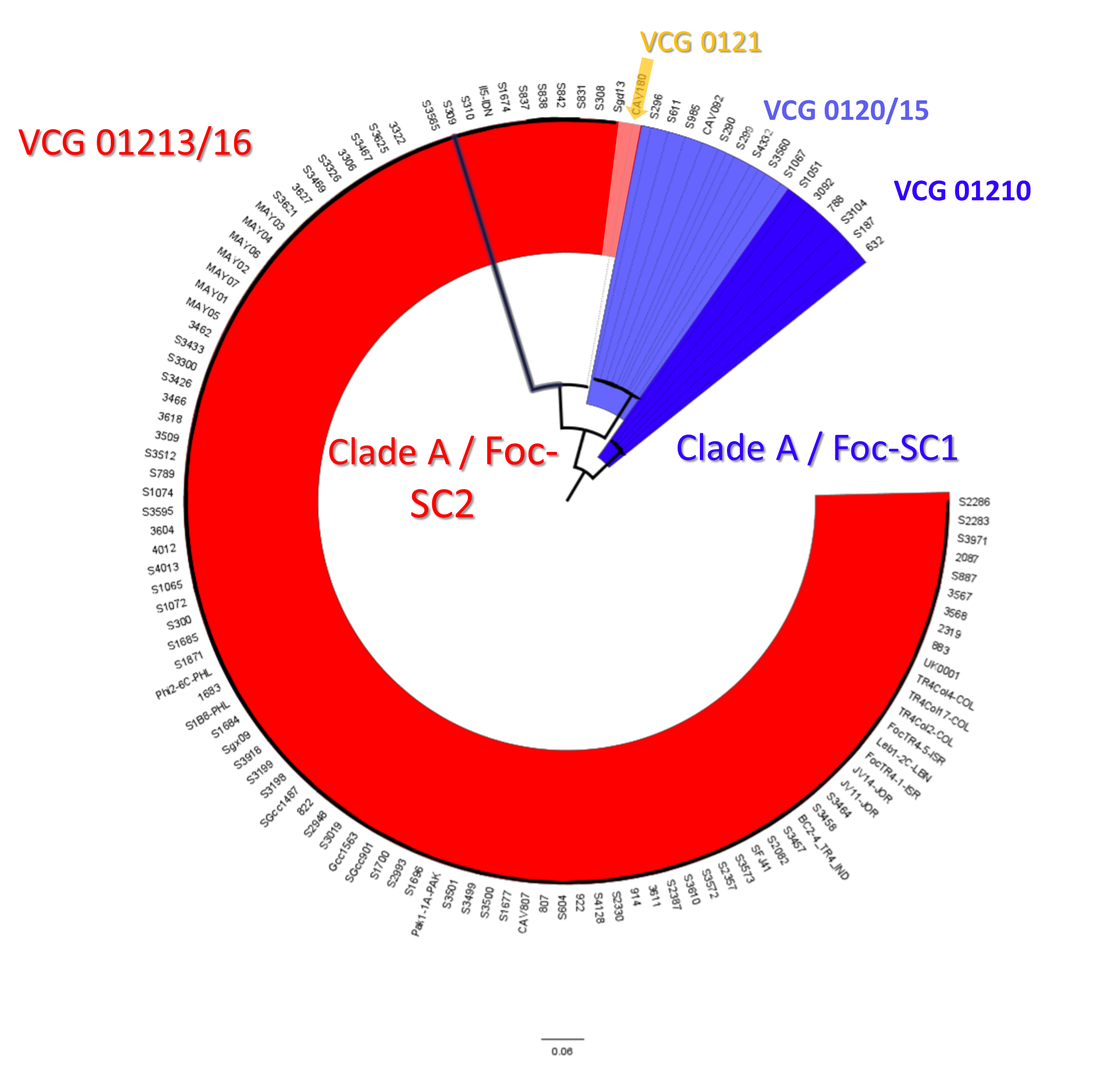

### Fig.S4

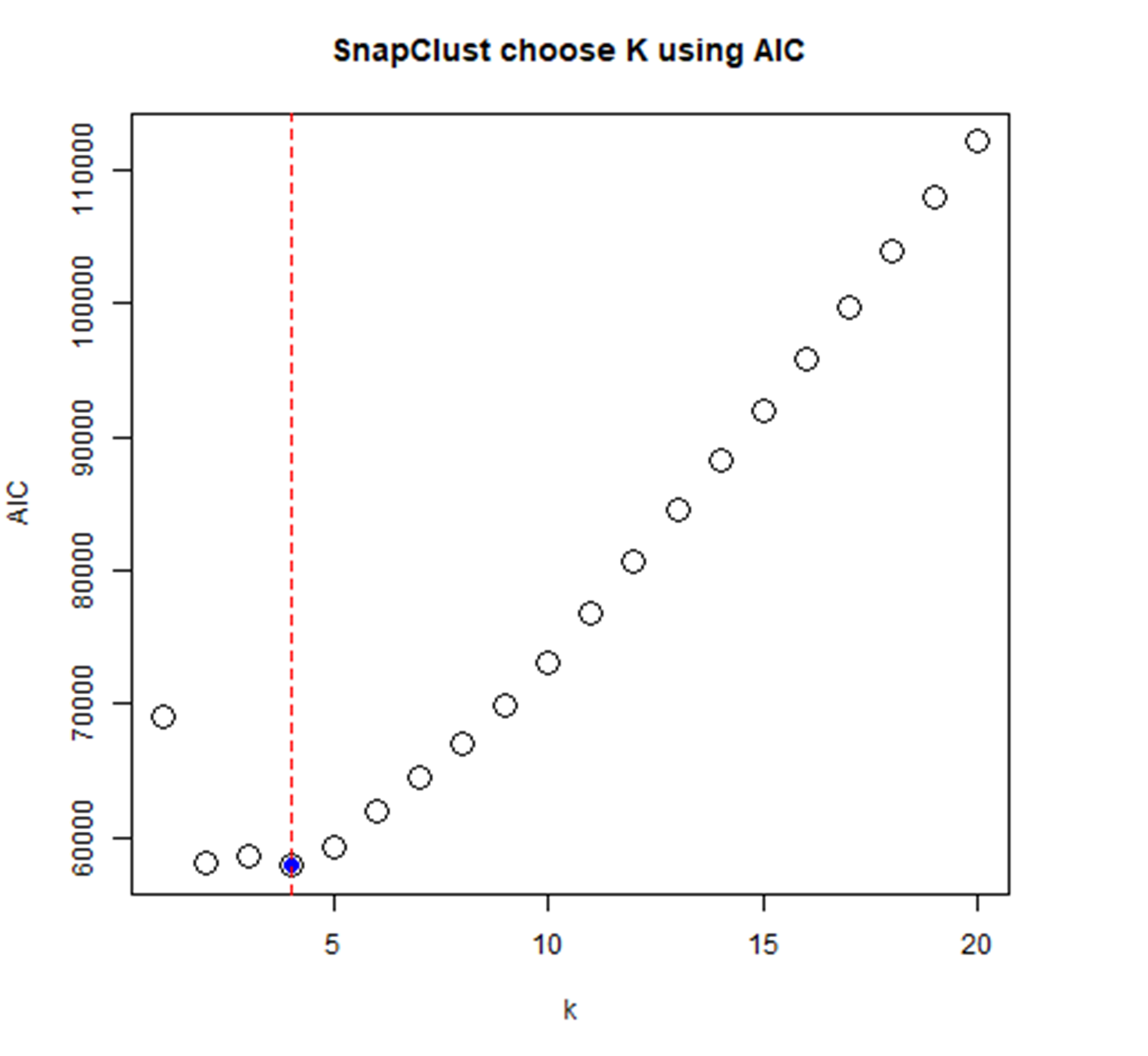

### Fig.S5

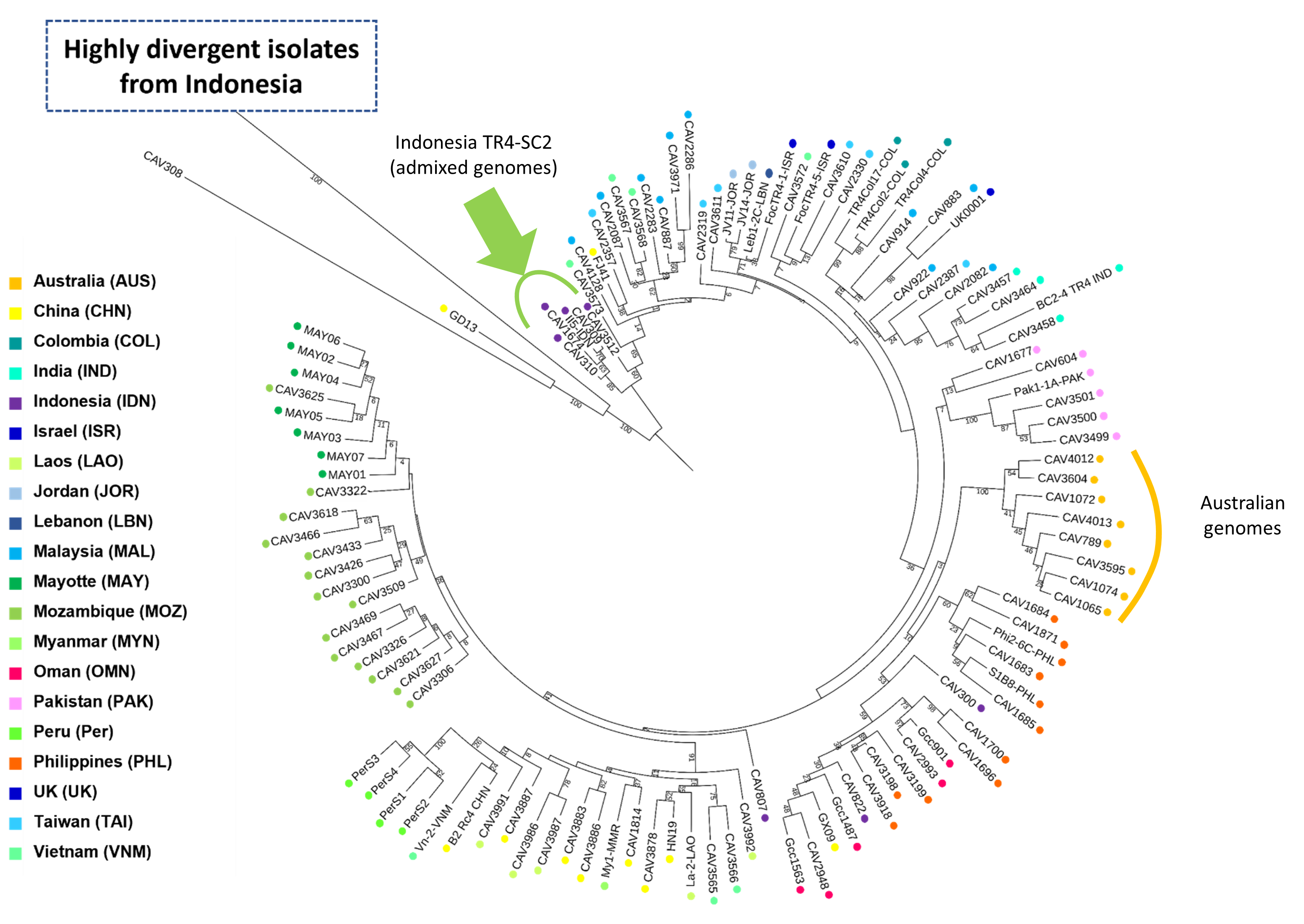

### Fig.S6

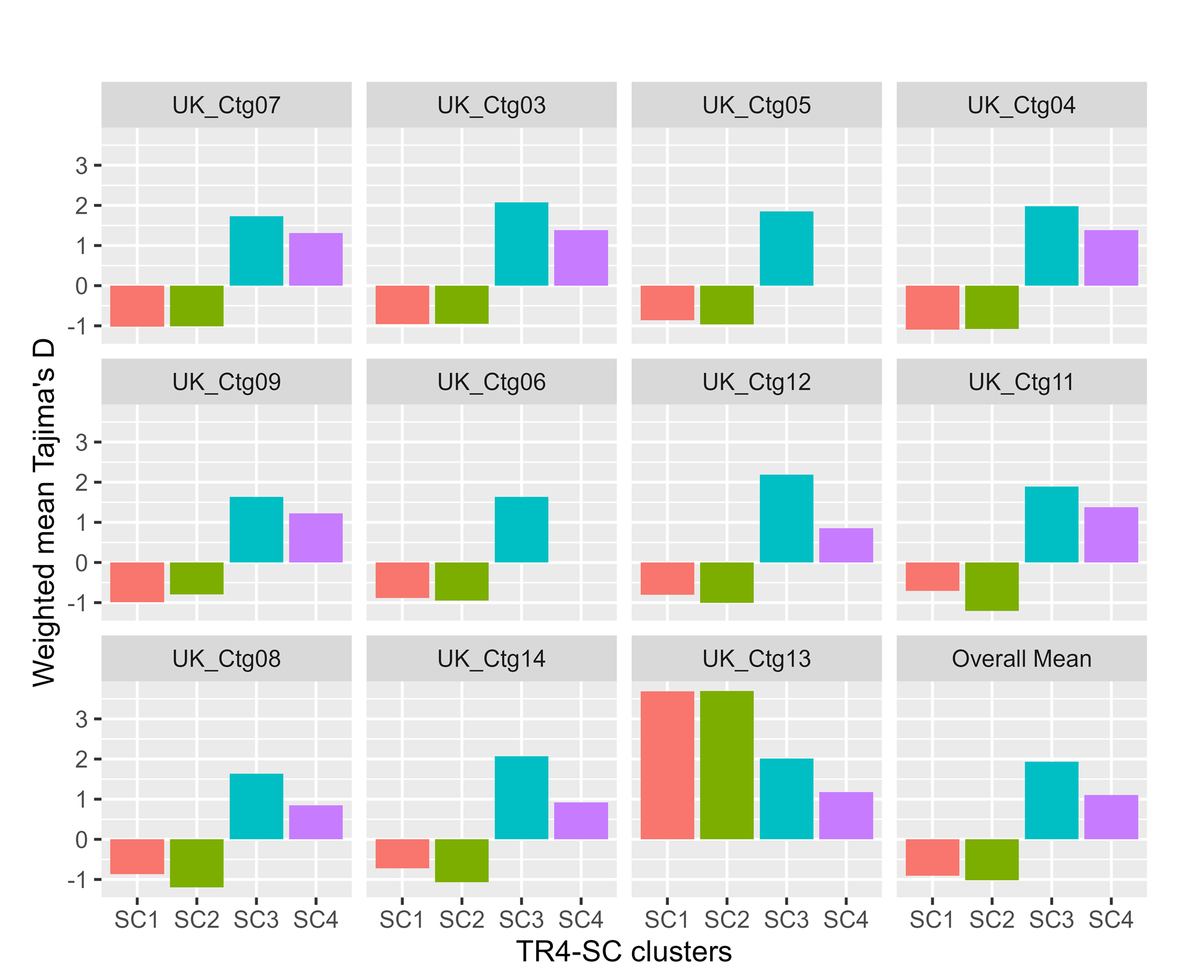

### Fig.S7

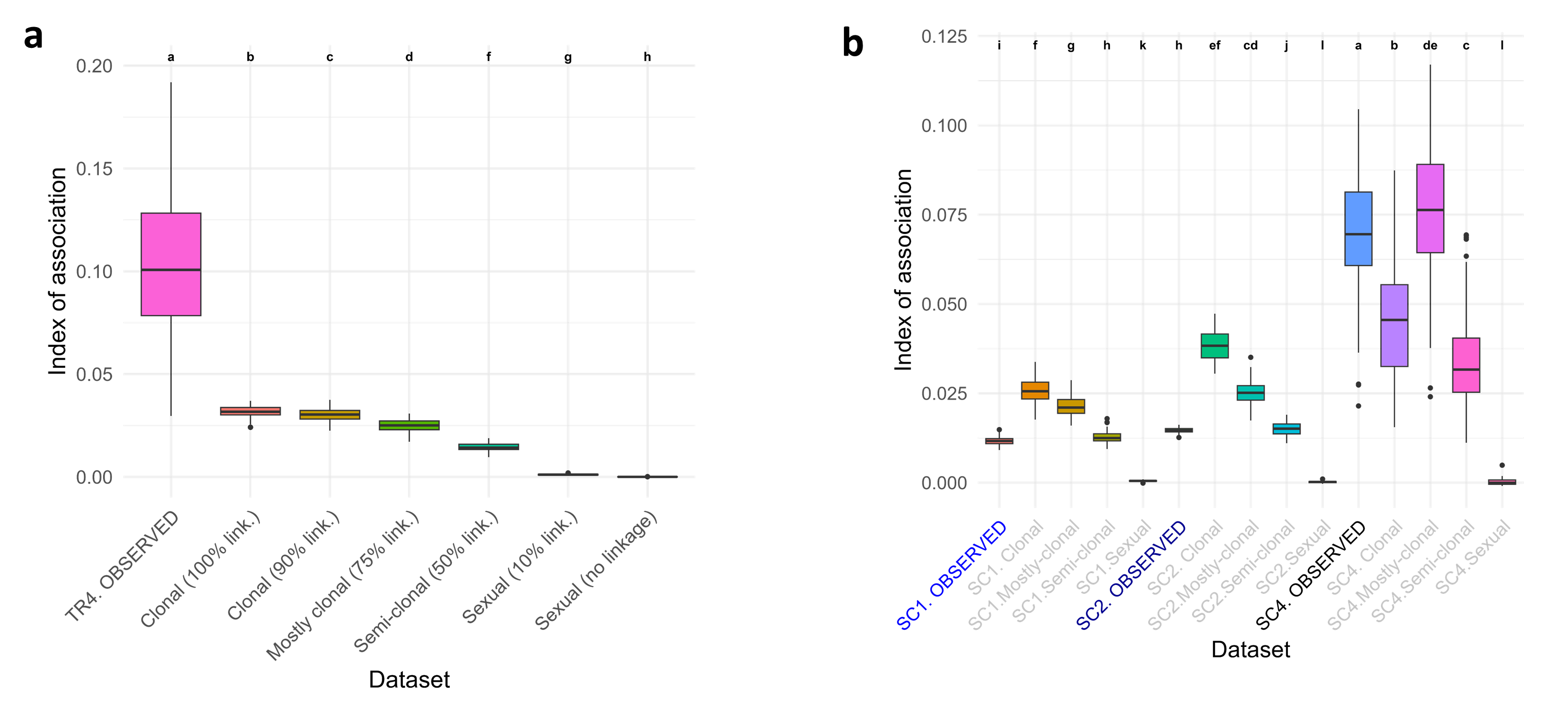
